# Structure of the complete extracellular bacterial flagellum reveals mechanism for flagellin incorporation

**DOI:** 10.1101/2024.09.25.614892

**Authors:** Rosa Einenkel, Kailin Qin, Julia Schmidt, Natalie S. Al-Otaibi, Daniel Mann, Tina Drobnič, Eli J. Cohen, Nayim Gonzalez-Rodriguez, Jane Harrowell, Elena Shmakova, Morgan Beeby, Marc Erhardt, Julien R. C. Bergeron

## Abstract

The bacterial flagellum is essential for motility, adhesion, and colonization in pathogens like *Salmonella enterica* and *Campylobacter jejuni*. Its extracellular structure comprises the hook, hook-filament junction, filament, and filament cap. The native structures of the hook-filament junction and the cap remain elusive, leaving the molecular details of cap-mediated filament assembly largely uncharacterized. Here, we report the structure of the complete extracellular flagellum, encompassing the hook, hook-filament junction, filament, and cap. This structure reveals intermediates of filament assembly, providing a molecular blueprint for flagellin folding and insertion at the filament tip. Mutagenesis and functional assays demonstrate the crucial roles of the cap’s terminal regions in flagellin incorporation, and of the structural integrity of the hook-filament junction. Finally, the structure of the cap and hook-filament junction prior to filament assembly reveals the structural basis for the initiation of filament assembly. Collectively, this study provides comprehensive insights into flagellum assembly and how flagellin incorporation is coupled with its secretion.

## Introduction

The flagellum is the most prominent extracellular structure in bacteria, with a molecular weight in the hundreds of megadaltons. It allows them to move within their environment through the rapid rotation of its propeller-like filament. In many human pathogens, including the prominent gastrointestinal pathogens *Salmonella enterica* (*S. enterica*) and *Campylobacter jejuni* (*C. jejuni*), the flagellum also plays an important role in infection, because of its ability to promote adhesion and colonization^1, 2^. Structurally, the flagellum can be divided into three major components: the basal body, the hook, and the filament (Figure 1A). The hook, a helical assembly of hundreds of subunits of FlgE, functions as a universal joint that connects the extracellular filament to the membrane-embedded basal body^3–5^. The flagellar filament, a multi-micron structure comprising tens of thousands of subunits of a single protein, the flagellin, facilitates bacterial motility through its rotation^6, 7^. Structural studies have revealed that the filament is a helical assembly consisting of 11 protofilaments^8–10^. A ring-like structure formed by the proteins FlgK and FlgL, termed the hook-filament junction (HFJ), forms the connection between the flexible hook and the rigid filament^11^. The structures of FlgK and FlgL have been previously reported, but only in their monomeric form^12–16^. In order to assemble a filament, flagellin monomers are secreted through the flagellar basal body and hook and finally polymerize at the distal end. The insertion of flagellin into the growing filament is mediated by the filament cap complex, which is composed of five copies of the protein FliD^17–19^. Based on the structure of the filament cap complex in isolation, we previously proposed that FliD forms a stable pentameric complex that rotates to enable the incorporation of new flagellin subunits into the respective protofilaments^20^. How the filament self-assembles, however, has remained elusive. Particularly, the molecular mechanism of the filament cap in facilitating the assembly of the filament and the function of the HFJ as a template for correct filament assembly are not yet understood.

**Figure 1.**
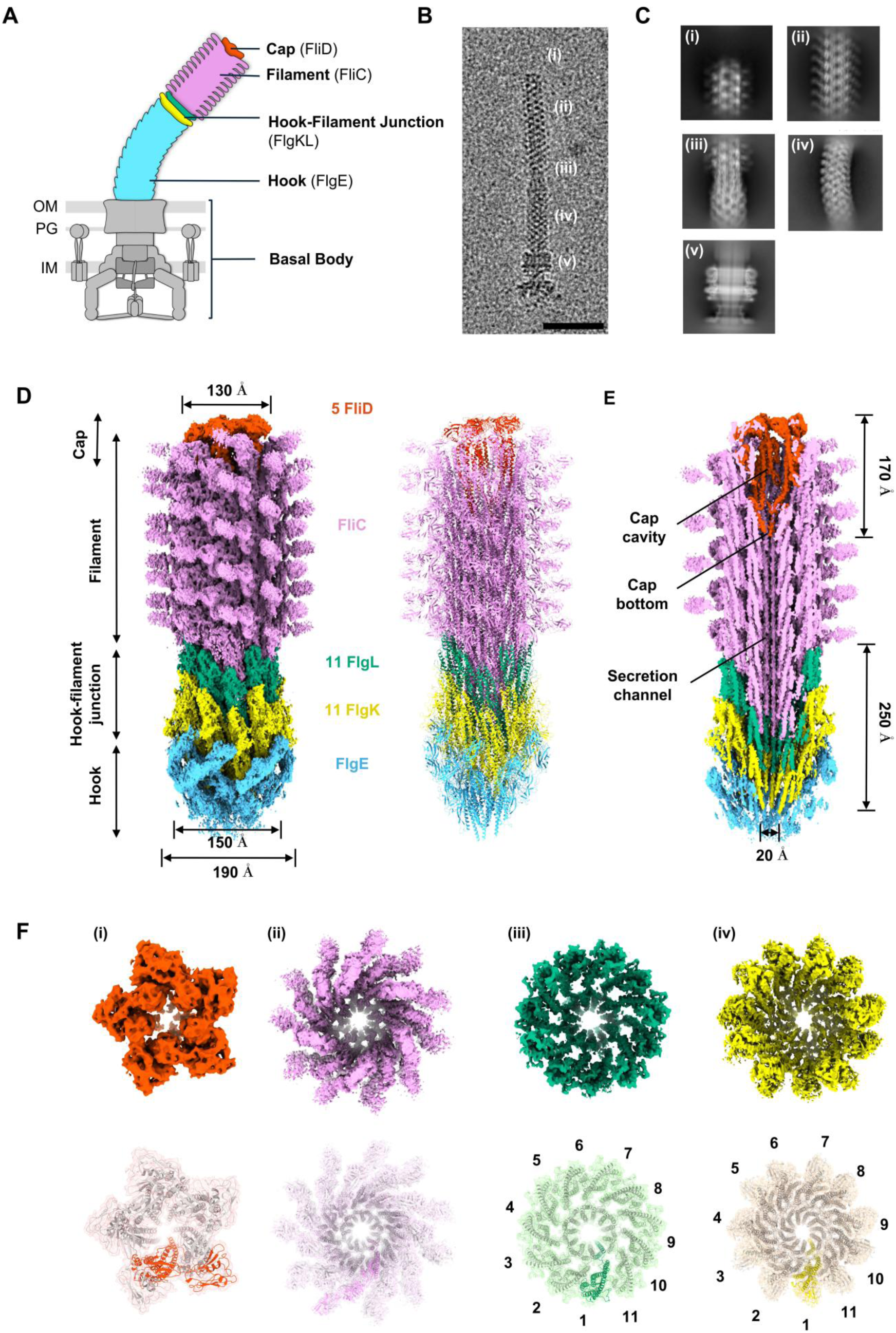
The complete structure of the extracellular flagellum. (A) Schematic representation of a Gram-negative bacterial flagellum. IM, inner membrane; PG, peptidoglycan layer; OM, outer membrane. (B) Representative micrograph displaying an intact flagellum: (i) cap region, (ii) filament, (iii) hook- filament junction, (iv) hook and (v) basal body, scale bar: 50 nm. (C) 2D class averages of the (i) cap region, (ii) filament, (iii) hook filament junction, (iv) hook and (v) basal body. (D) Connected density map of the extracellular flagellar complex from the hook to the cap. The map of the flagellar tip and the map of the hook-filament junction are refined individually and connected through multiple layers of flagellin. The atomic model of the entire extracellular flagellum is displayed (PDB: 9GNZ, 9GO6). (E) Cross-section view of the merged map with labels of dimension of FliD cap and FlgKL junction. (F) Top-down view of each section of the extracellular flagellum. Models are fitted into the map, which is displayed in the same color but with transparency, and the subunits for each protein are highlighted. (i) FliD pentamer, (ii) FliC filament, (iii) FlgL layer with its stoichiometry in the HFJ, (iv) FlgK layer with its stoichiometry in the HFJ. See also Movie S2.

Here, we report the structure of the filament cap complex in its native environment, assembled at the distal end of the flagellar filament. Critically, we were able to determine this structure at various stages of flagellin incorporation into the filament, allowing us to identify the molecular steps involved in this process. Using mutagenesis and functional assays, we demonstrate that the FliD terminal domains are essential for flagellin folding and incorporation into the filament. Furthermore, we report the structure of the intact HFJ within a fully assembled flagellar filament, experimentally confirming the proposed 11:11 stoichiometry of the FlgKL proteins and revealing the molecular details of its gasket-like role in isolating the filament from the hook. Structure-guided mutagenesis, which destabilizes the FlgKL interface, highlights the importance of FlgKL inter-molecular interactions for structural integrity and proper function of the flagellar apparatus. Finally, we report the structure of the cap assembled on the HFJ, corresponding to the state of the complex prior to filament assembly, which reveals the structural changes that occur in the cap upon the initiation of filament formation. Collectively, these results reveal a detailed model for cap-mediated flagellin insertion, illustrating how flagellin incorporation is coupled with secretion.

## Results

### The complete structure of the extracellular flagellum

We previously reported the structure of the flagellum cap complex in isolation^20^, revealing a pentameric complex with 5-fold symmetry. To determine the structure of the cap complex in its native context, we applied cryogenic electron tomography (cryo-ET) on intact *S. enterica* cells. We collected 68 tomograms, providing 252 flagellar ends for subtomogram averaging, leading to a map with a resolution of ∼25 Å (Figure S1, Movie S1). In this map, we observed a seam in the cap, showing asymmetric positioning of the FliD subunits within the filament (Figure S2). This indicates structural differences between the native state and the structure of the isolated cap complex^20^. However, we could not resolve the details of FliD arrangements and their interactions with flagellins from this map because of its low resolution. To resolve this, we aimed to determine this structure using single-particle cryo-EM.

We genetically modified *S. enterica* to generate flagellum complexes with short filaments, reasoning that these shorter structures would be more amenable to cryo-EM imaging and data processing. To increase flagella yield during purification, we exchanged the native promoter of the flagellar master regulator FlhDC with the strong synthetic constitutive promoter P*proB*, leading to hyperflagellated cells^21, 22^. To control filament length, we locked the cells in the production of the flagellin FliC and exchanged the native *fliC* promoter with an inducible P*tetA* promoter. We observed that a 30 min induction time for flagellin production resulted in sufficiently short filaments, reducing the risk of filament breakage and hence loss of the filament cap during purification (Figure S1D). Flagella were then purified and found to contain the intact basal body, hook, junction, filament, and cap (Figure 1B-C), as described previously^23–25^.

Cryo-EM analysis of these purified flagella with short filaments (Figure S2) allowed us to obtain independent maps for the cap and HFJ to 3.7 Å and 2.9 Å resolution, respectively (Table S1). For the cap complex, we were able to refine this map further, to 3.3 Å, however in this map, the D2 and D3 domains of one FliD subunit were not resolved, indicating structural heterogeneity. To address this, 3D classification was employed to identify the different states of the cap, leading to a final map with all five FliD subunits (Figure S2). The final model of the cap contains 5 FliD subunits, and 17 copies of FliC in the filament; the final model of the HFJ contains 13 copies of FlgE for the hook, 11 copies of FlgK and FlgL, and 14 copies of FliC. We emphasize that despite the structural similarity between FlgK and FlgL, we were able to unambiguously build these into our map because of its high resolution, where side- chains are readily identifiable. Finally, we connected the HFJ and cap maps by aligning them through multiple copies of FliC molecules, leading to a composite map encompassing the distal end of the FlgE hook, FlgKL HFJ, FliC filament, and FliD cap (Figure 1D-E, Movie S2).

This composite structure shows that the cap has a width of ∼130 Å and a length of ∼170 Å (Figure 1E). The D2-D3 plane of the FliD pentamer is tilted, which is consistent with our tomography data (Figure 1D-E, Figure 2A, Figure S1). We observed the presence of a cavity enclosed by D2-D3 and D0-D1 of the cap, consistent with early low-resolution data^19^. FlgK and FlgL assemble into the HFJ in individual layers (Figure 1D-F). The width of each layer is ∼150 Å for FlgL and ∼190 Å for FlgK (Figure 1D). The total length of the HFJ from the top of FlgL to the bottom of the FlgK layer is ∼250 Å. Eleven FlgL subunits form the distal layer below the filament, and 11 FlgK subunits form the proximal layer of the junction stacked on top of the hook (Figure 1F). Hence, the stoichiometry and arrangement of FlgKL are consistent with previous suggestions that were made based on structural modeling^12, 14, 15^.

**Figure 2.**
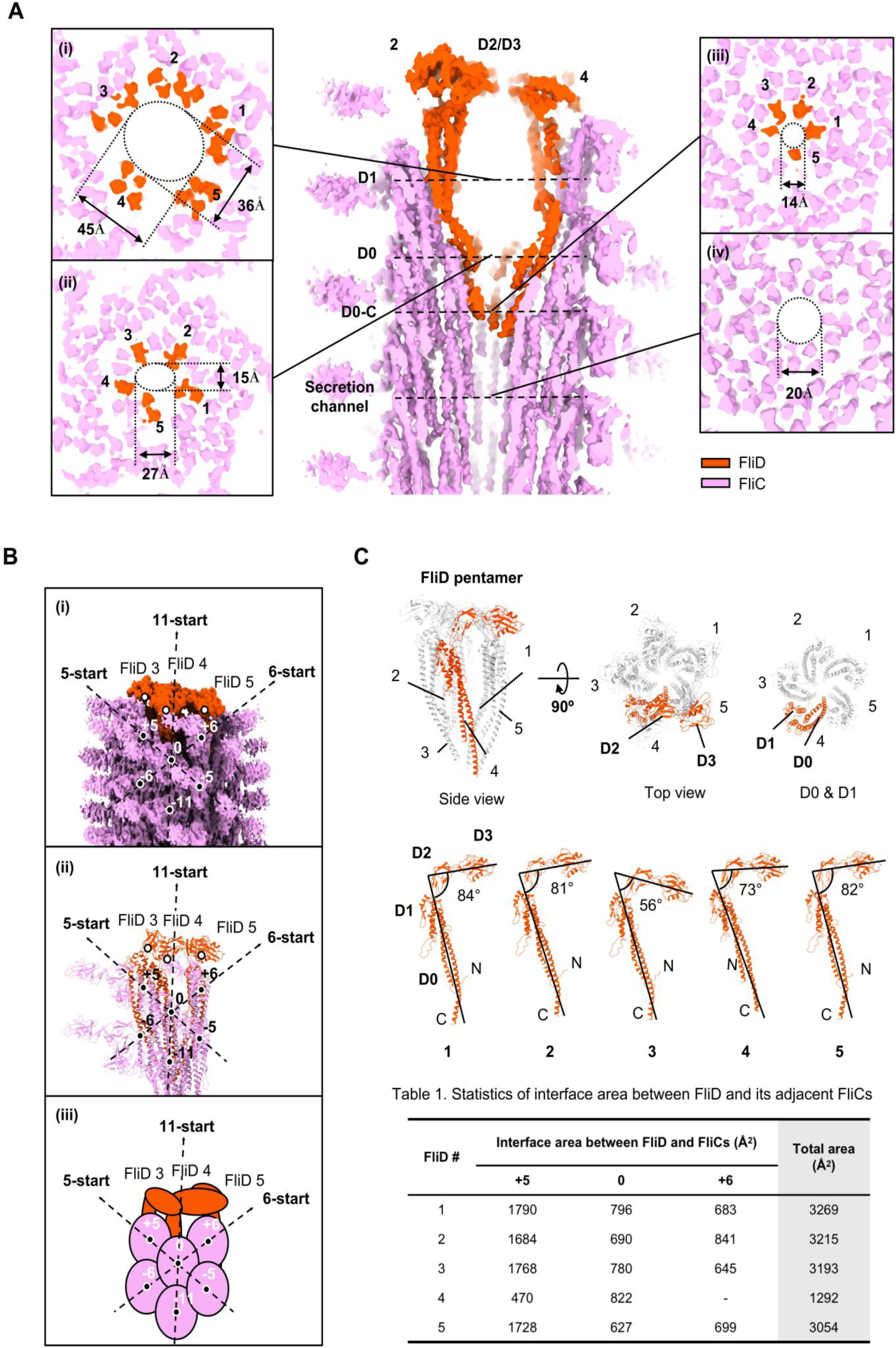
Structure of the native cap complex, and its interaction with the filament. (A) Cross section of FliD cap at the distal end of the filament and the dimension of the cavity. (i) Cross section at D1 helices at the widest part of the cavity, (ii) Cross section at D0 helices, (iii) Cross section at D0-C helices at the bottom of the cavity, (iv) Cross section of the FliC filament below FliD cap. (B) Arrangement of FliD and FliC in the cap-filament complex. (i) Symmetry array of FliC in the filament and FliD position relative to FliC, (ii) Arrangement of subunits displayed as atomic model (PDB: 9GNZ), (iii) Schematic representation of FliC symmetry and relative positions of FliD and FliC. (C) Overall atomic structure of FliD cap and structural polymorphism among FliD. See also Figure S3.

### Structure of the native cap complex, and its interaction with the filament

The flagellar cap complex is composed of five FliD subunits assembled into a pentamer with an overall shape resembling an acorn. Consistent with our previous structure of the cap in isolation^20^, the FliD monomer consists of four domains, termed D0 to D3. However, in the native structure, the D2-D3 domains assemble into a tilted, star-shaped plane and the D0-D1 helices form a plug-like structure beneath (Figure 2A). The D2-D3 plane and D0-D1 helices surround a large cavity, much larger than that described previously^19^, and the D0 domains form an almost closed constriction in the secretion channel (Figure 2A). At its widest, around the D1 domains, the cavity adopts an oval shape, ∼45 Å wide along the long axis and ∼30 Å wide along the short axis (Figure 2A.i). The cavity is narrower across the D0 domain, ∼27 Å for the long axis, and ∼15 Å for the short axis (Figure 2A.ii). The bottom of the cavity is formed by C-terminal D0 domains (hereafter referred to as D0-C), which form a narrow constriction with a diameter of ∼14 Å, continuous with the secretion channel in the filament (Figure 2A.iii). This constriction is narrower than that of the channel in the filament, which is 20 Å (Figure 2A.iv).

The D0-D1 domains of FliD are oriented vertically, while the D2-D3 domains are horizontal and point in the counterclockwise (CCW) direction (Figure 2C). The D3 domain of one FliD stacks on the D2 domain of the adjacent FliD in CCW direction, e.g. D3 of FliD 4 is stacked on top of D2 of FliD 5 (Figure 2C). Owing to the varying heights of each FliD subunit, the D2-D3 plane is tilted, with angles between D0-D1 and D2-D3 of 84° (FliD 1), 81° (FliD 2), 56° (FliD 3), 73° (FliD 4) and 82° (FliD 5) (Figure 2C).

FliD subunits exhibit a pseudo-symmetric arrangement, forming an asymmetric unit with three adjacent FliC molecules, labeled FliC 0, FliC +5, and FliC +6 (Figure 2B). This pseudo-symmetry pattern is consistent across the five FliD molecules. Using PISA^26, 27^, we analyzed the interactions between the FliD subunits and their respective adjacent FliC molecules within the aforementioned asymmetric units (Figure S3). FliD subunits 1, 2, 3, and 5 form an extensive interface with FliC +5, with the interface being twice as large as that with FliC 0 or FliC +6. In contrast, FliD 4 only interacts with FliC +5 and FliC 0, but not with FliC +6 (Table S2). Moreover, FliD 4 forms a significantly smaller interface with FliC +5. These unique features indicate that FliD 4 participates in the incorporation of a new FliC, which occurs in the gap between FliD 4 and FliC +6.

### Conformational changes to the cap upon filament assembly are coupled with flagellin folding

As indicated above, we noticed that one FliD subunit is not well resolved in our initial map (Figure S2). Therefore, we hypothesized that this FliD subunit is highly flexible and adopts different conformations, whereas the other FliD subunits are more stable. To verify this, we performed 3D variability analysis in cryoSPARC^28^. This allowed us to obtain individual maps with assembly intermediates, whereby flagellin subunits are in the process of being incorporated into the filament, and FliD monomers adopt distinct conformations to that of the final map described above (Figure 3, Movie S3). To clearly visualize the FliD movements during FliC incorporation, we colored the FliD subunits that undergo conformational changes as salmon (FliD 3), red (FliD 4), and orange (FliD 5) (Figure 3A-B and Movies S4-5). Notably, we were able to reconstruct the molecular motion in the cap complex during filament assembly. During the incorporation of a FliC molecule, the D0-D1 domains of FliD 4 rise upwards (Δ = 25 Å) and shift in the clockwise (CW) direction (θ = 18°). The D2 domain of FliD 4 also rises, thereby causing the rise of the D3 domain of FliD 3, while the N-terminal loop of FliD 4 rises and turns 90° upward (Movies S4-5). Finally, the D3 domain of FliD 4 rises, which is caused by the rise of the D2 domain of FliD 5, completing the cycle.

**Figure 3.**
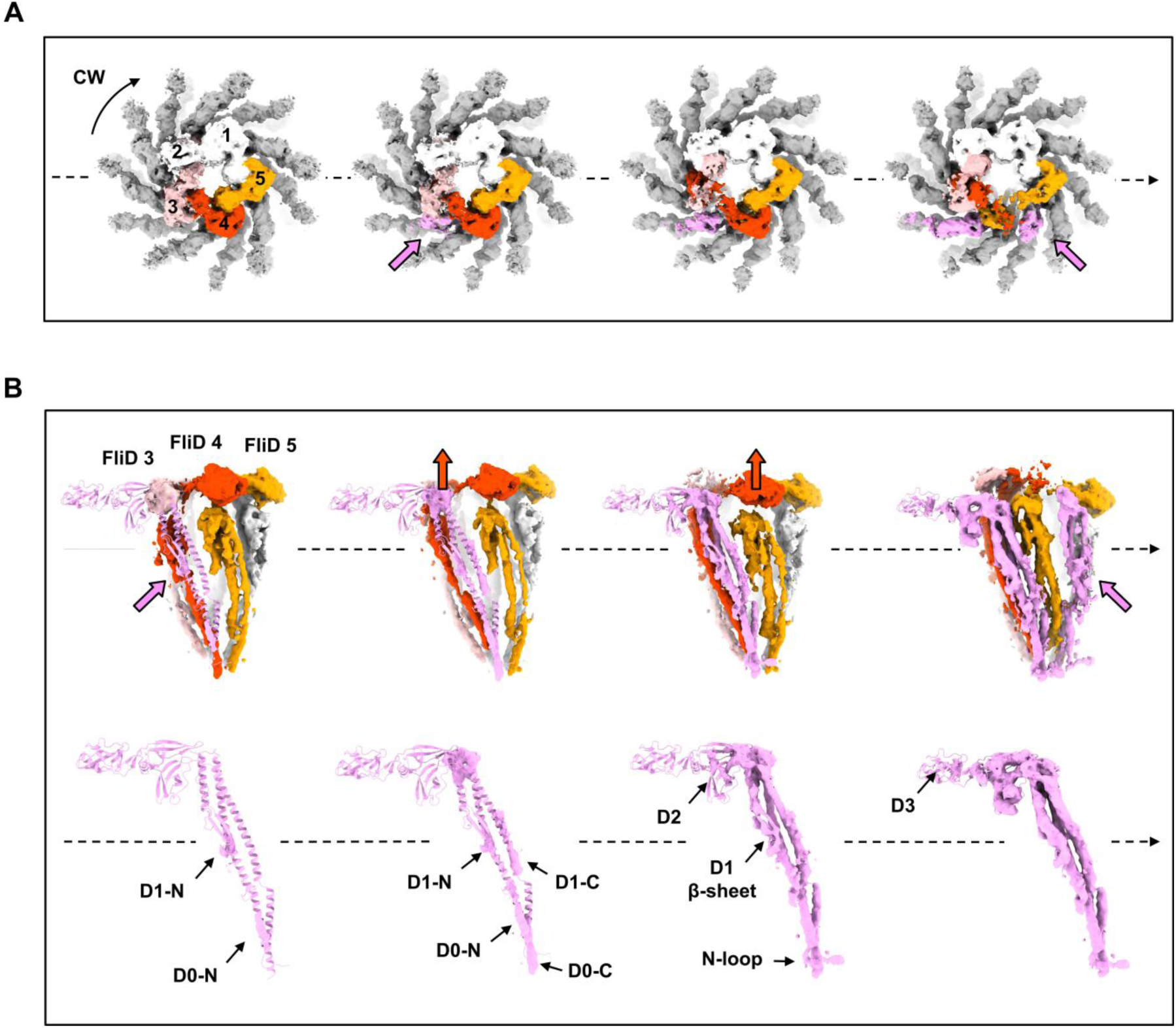
Conformational changes to the cap and flagellin subunits upon filament assembly. (A) Top-down view of clockwise rotation of the FliD cap (black arrow) and counterclockwise incorporation of FliC (pink arrow) over time. FliD subunits that undergo conformational changes are colored salmon, red, and orange. FliD subunits are labeled in the same order, as shown in Figure 2C. (B) Side view of the conformational changes of the FliD cap (red arrows) and FliC incorporation (pink arrows) over time, with a focused view of the density appearing at each state of FliC incorporation. See also Movies S3, S4, S5.

By analyzing the results of the 3D variability analysis frame-by-frame, we also reconstructed the steps of FliC folding upon incorporation into the filament (Movie S3, Figure 3A-B). Densities for the D0 and D1 helices of the N-terminus, which are close to the D0-C domain of FliD and the D1 domain of FliC 0, respectively, emerge initially (Figure 3B). Subsequently, densities of helices in the C-terminal D0-D1 domains emerge, which are close to the N-terminal D0 domain (hereafter referred to as D0-N) of FliD and the D1 domain of FliC +6, respectively, and more densities can be seen in the helices in D0-N and D1-N of FliC. Following this, helices in the D0 and D1 domains, the β-sheet and loops in the D1 domain, and the loop at the N-terminus are completed. At this stage, the density of the D2 domain begins to emerge. Later, the D2 is completed and the D3 domain starts to appear (Figure 3B). Notably, before the maturation of one FliC monomer is complete, the incorporation of the next FliC molecule begins (Figure 3B). Given the spatial patterns of domain appearance, we propose that the folding sequence of FliC is as follows: helices of D0-N and D1-N, helices of D0-C and D1-C, β-sheet and loop in D1 and N-loop, and D2 and D3 domains.

### FliD terminal regions mediate flagellin insertion at the flagellar distal end

Based on our structural studies, we investigated the importance of FliD terminal domains in filament elongation in *S. enterica* using genetic engineering and functional assays. We focused on amino acids located in the FliD-FliC interaction site within the D0-D1 domains of FliD. We analyzed bacterial motility by assessing the ability of the mutants to swim through semi-solid agar (Figure 4). Substituting conserved hydrophobic residues in the D0 domain of FliD to serine has no (V9S, F461S) or mild (L22S, F440S, L443S, M446S, and L450S) impact on swimming ability. Similarly, serine mutations in residues of the D1 domain involved in flagellin interactions result in mild (Y296S) or no (R319S) reduction of motility. Double mutation of the D1 domain residues (Y296S R319S) and substitution of Y296 with arginine results in a 10% decrease of motility.

**Figure 4.**
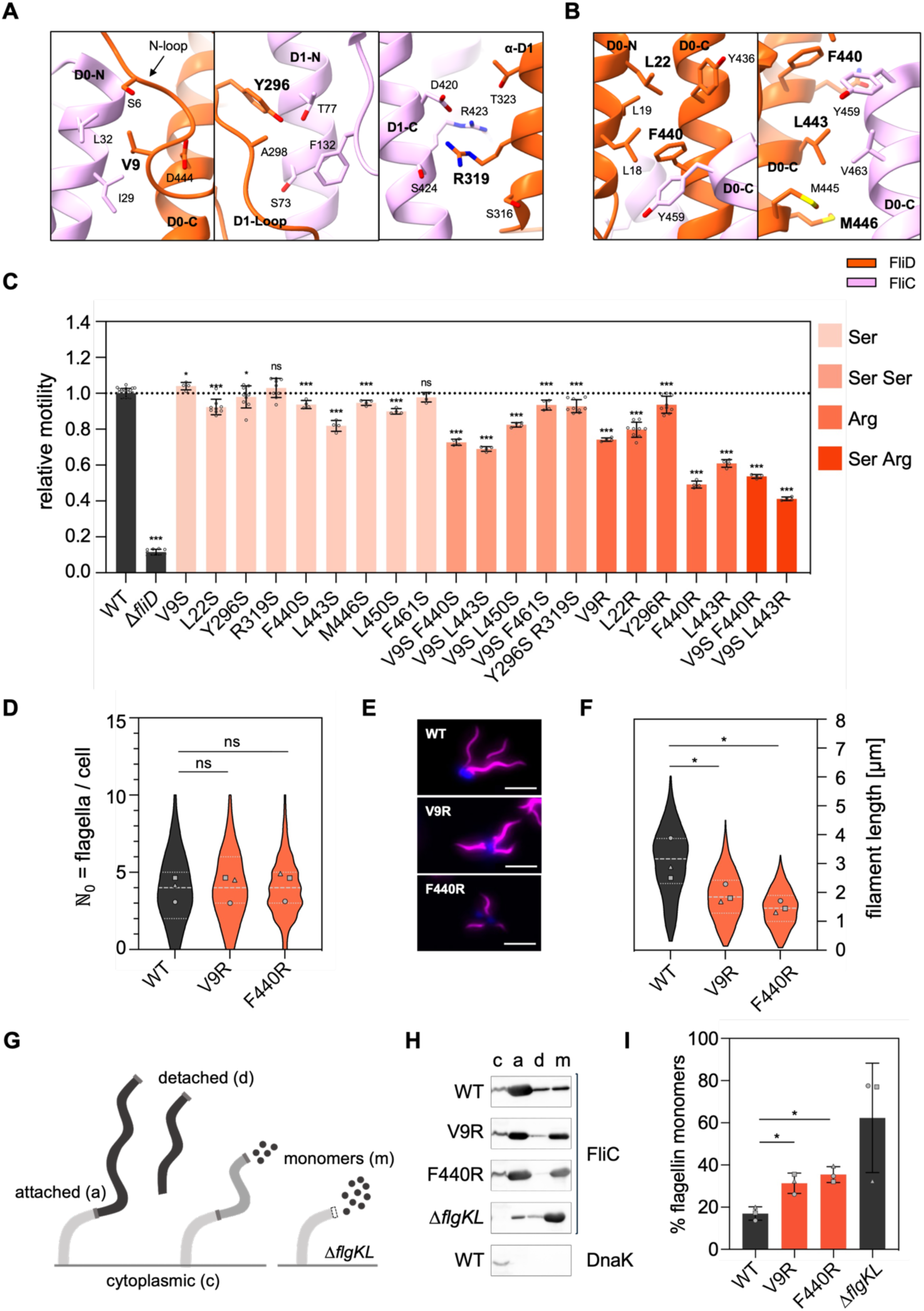
FliD terminal regions mediate flagellin insertion at the flagellar distal end. (A) Interactions between FliD and FliC at position +5. Residues chosen for mutagenesis and related domains of FliD are highlighted in bold black. (B) Interactions between FliD and FliC at position +6. Residues chosen for mutagenesis and related domains of FliD are highlighted in bold black. (C) Relative motility of *S. enterica* FliD point mutants analyzed using soft-agar motility plates, quantified after 4-5 h incubation at 37 °C. Diameters of the motility halos were measured using Fiji and normalized to the WT. Bar graphs represent the mean of at least 3 biological replicates with standard deviation error bars. Replicates shown as individual data points. Ser, single substitution to serine; Ser Ser, double substitution to serine; Arg, single substitution to arginine; Ser Arg, double substitution to serine and arginine. (D) Quantification of the number of flagella per cell in the WT and FliD mutants. The number of flagella per cell (ℕ0) was determined for n > 150 individual bacteria per strain for 3 biological replicates. Violin plots represent the distribution of the data of all replicates including the median (dashed line) and quartiles (dotted lines). Data points represent the means of each biological replicate. (E) Representative fluorescence microscopy images of the WT and FliD mutants (V9R and F440R). Filaments (FliC T237C) were labeled with Dylight555 Maleimide (magenta) and DNA counterstained with DAPI after pulsed *flhDC* induction. Scale bar: 3 µm. (F) Quantification of the filament length in µm in the WT and FliD mutants. The filament length was determined for n > 150 individual filaments per strain for 3 biological replicates. Violin plots represent the distribution of the data of all replicates including the median (dashed line) and quartiles (dotted lines). Data points represent the means of each biological replicate. (G) Schematic overview of the experimental setup to determine flagellin leakage during filament formation. (H) Representative immunodetection of cytoplasmic, cell-attached, detached and monomeric FliC in the WT, the FliD mutants (V9R and F440R) and Δ*flgKL* control strain. DnaK immunodetection of the WT shown as representative lysis control. c, cytoplasmic flagellin; a, attached flagellin; d, detached flagellin; m, monomeric flagellin. (I) Proportion of secreted flagellin as a percentage of the total flagellin amount of the WT, the FliD mutants (V9R and F440R) and a Δ*flgKL* control strain. Leakage assays were performed in 3 biological replicates. Statistical annotations were calculated with a two-tailed Student’s t-test, on the means of each biological replicate (*, p < 0.05; **, p < 0.01; ***, p < 0.001; ns, non-significant). See also Figures S4A, S4B, S4E.

However, double serine mutants of the D0 domain (V9S F440S, V9S L443S, V9S L450S, V9S F461S) exhibit more pronounced motility defects (70-90% of wild-type (WT) motility) than their respective single mutants. Furthermore, arginine substitutions in terminal residues result in significant motility reductions: F440R and L443R and the V9S F440R double mutant display approximately 50% WT motility, while V9R and L22R retain about 80%. The most severe defect is observed in the V9S L443R double mutant, with 41% of WT motility. Overall, our data suggest that the FliD D0 and D1 domains are robust towards mutations. Substitutions to small, polar residues are tolerated, while substitutions to larger, charged residues impair motility.

To confirm that the motility defects result from impaired filament assembly, we studied the flagellation patterns of the cap mutants using fluorescence microscopy (Figure 4E and S4A-B). We employed a pulsed *flhDC* induction setup (Figure S4E) to ensure synchronization of flagella biosynthesis and visualized the filaments using a fluorophore-coupled maleimide dye (Figure 4D-F, S4A-B). The WT and mutant bacteria possess an average of four filaments per cell, indicating that the mutations do not affect the genetic regulation of flagella biosynthesis (Figure 4D). Importantly, filament length is significantly reduced in the analysed cap mutants with severely reduced motility (V9R and F440R), being on average 38% (V9R) or 52% (F440R) shorter than WT filaments (Figure 4F). These results confirm that the reduced motility resulting from these mutations is due to impaired filament assembly.

To verify that the reduced filament length is not caused by inefficient secretion or premature degradation of FliD, we compared cellular and secreted levels of FliD in mutant and WT bacteria (Figure S4C-D). We used the pulsed induction setup (Figure S4E) and included a non-secreting control, deficient in the flagellar export apparatus (Δ*fliP*), and a filament-assembly mutant (Δ*flgKL*). Cellular FliD levels do not significantly differ between mutants and the WT, excluding premature FliD degradation (Figure S4C). Secreted FliD levels are comparable between the mutants and the WT, indicating that FliD secretion is not affected. However, FliC secretion is significantly higher in the FliD mutants (Figure S4D), suggesting inefficient incorporation of flagellin molecules into the growing filament.

We next confirmed flagellin leakage during filament formation by separating cellular flagellin, flagellin attached to the cell body, detached filaments, and leaked flagellin monomers using differential (ultra)centrifugation (Figure 4G). In the WT, 17% of secreted FliC is found in the supernatant in monomeric form, whereas most flagellin is incorporated into the filament (Figure 4H-I). In contrast, the V9R mutant exhibits a significant increase in monomeric flagellin in the supernatant (31%) and the F440R mutant displays an even greater increase (35%), consistent with fluorescence microscopy and secretion assay results. In a mutant lacking the HFJ (Δ*flgKL*), FliC is mostly present (62%) in the supernatant, confirming that this mutant does not permit filament assembly; however, we note that some level of cell-attached flagellin is detected in this mutant, likely due to enhanced secretion and aggregation of FliC monomers. Collectively, our data demonstrate that the filament cap mutants incorporated nascent flagellin less efficiently, resulting in shorter filaments and reduced bacterial motility.

### Structure of the HFJ, and its interactions with the hook and filament

As indicated above, using our short flagellum cryo-EM dataset, we were able to determine the structure of the intact HFJ, anchored to both the hook and the filament, to 2.9 Å resolution (Figure S2). In this structure, both FlgK and FlgL form 11-mer layers that separate the hook protein FlgE from the filament protein FliC (Figure 2D-E). These components - FlgE, FlgK, FlgL and FliC - are aligned to form a continuous protofilament along the 11-start symmetry axis (Figure S5A). The D0 domains of all four proteins face the lumen, creating a continuous channel with a consistent diameter of ∼20 Å (Figure S5B, Figure 1E). Helices in the D1 domain of all proteins, except FlgE, are entirely buried, whereas the D1 domain of FlgE, the D2 domains of FlgK, FlgL, and FliC, and the D3 domain of FliC are exposed to the extracellular environment (Figure S5A). The HFJ proteins follow a heterodimer pseudo-symmetry along the 11- start axis (Figure S5B).

FlgK and FlgL adopt three different modes to interact with adjacent proteins, corresponding to their position, with mode 1 being adopted only by subunit 11, mode 2 adopted by subunits 1, 3, 5, 7 and 9, and mode 3 adopted by subunits 2, 4, 6, 8 and 10 (Figure S5B-C). Along the 11-start axis, FlgK exclusively interacts with FlgL at +11 and FlgE at −11; FlgL exclusively interacts with FliC at +11 and FlgK at −11. Along the 5-start and 6-start axes, FlgK and FlgL interact with multiple subunits: FlgK interacts with FlgK/FlgL (+5), FlgK/FlgE (−5), FlgK/FlgL (+6), FlgK/FlgE (−6); Flg, interacts with FliC/FlgL (+5), FlgK/FlgL (−5), FliC/FlgL (+6), FlgK/FlgL (−6). Using PISA^26, 27^, we calculated the interaction interface area between each junction protein and the six adjacent subunits for the three interaction modes (Table. S2). Large surfaces of interactions are found along the 5-start and 11-start axes, rather than the 6-start axis, for both FlgK and FlgL. For FlgK, larger interfaces are found along the 5-start axis between two FlgK subunits, rather than interactions between FlgK and FlgE (−5) or between FlgK and FlgL (+5). For the FlgK subunit(s) adopting the interaction mode 3, the interface with FlgE (−11) is significantly larger (Figure S5C, Table S2). For FlgL, the interface area is similarly large when interacting with another FlgL or FlgK at −5. However, in mode 3, the interface areas with FliC (+5) and FlgL (−5) decrease substantially (Table S2).

It is noteworthy that there is no interaction between the hook protein FlgE and FlgL, nor between FlgK and the flagellin FliC. This likely explains why two proteins are required for the HFJ, whereas a single protein would not be able to provide a complete isolation between the hook and filament components of the flagellum.

### Mutations in the HFJ interface impair bacterial motility and filament stability

Based on the interaction analysis using PISA^26, 27^, residues Q111, Q118, D519 for FlgK and I44, L260 for FlgL were determined to be fully buried at the interface between FlgK and FlgL (Figure 5A). To test the role of these residues in HFJ function, we constructed mutants of these residues and assessed their ability to swim through semi-solid agar (Figure 5B).

**Figure 5.**
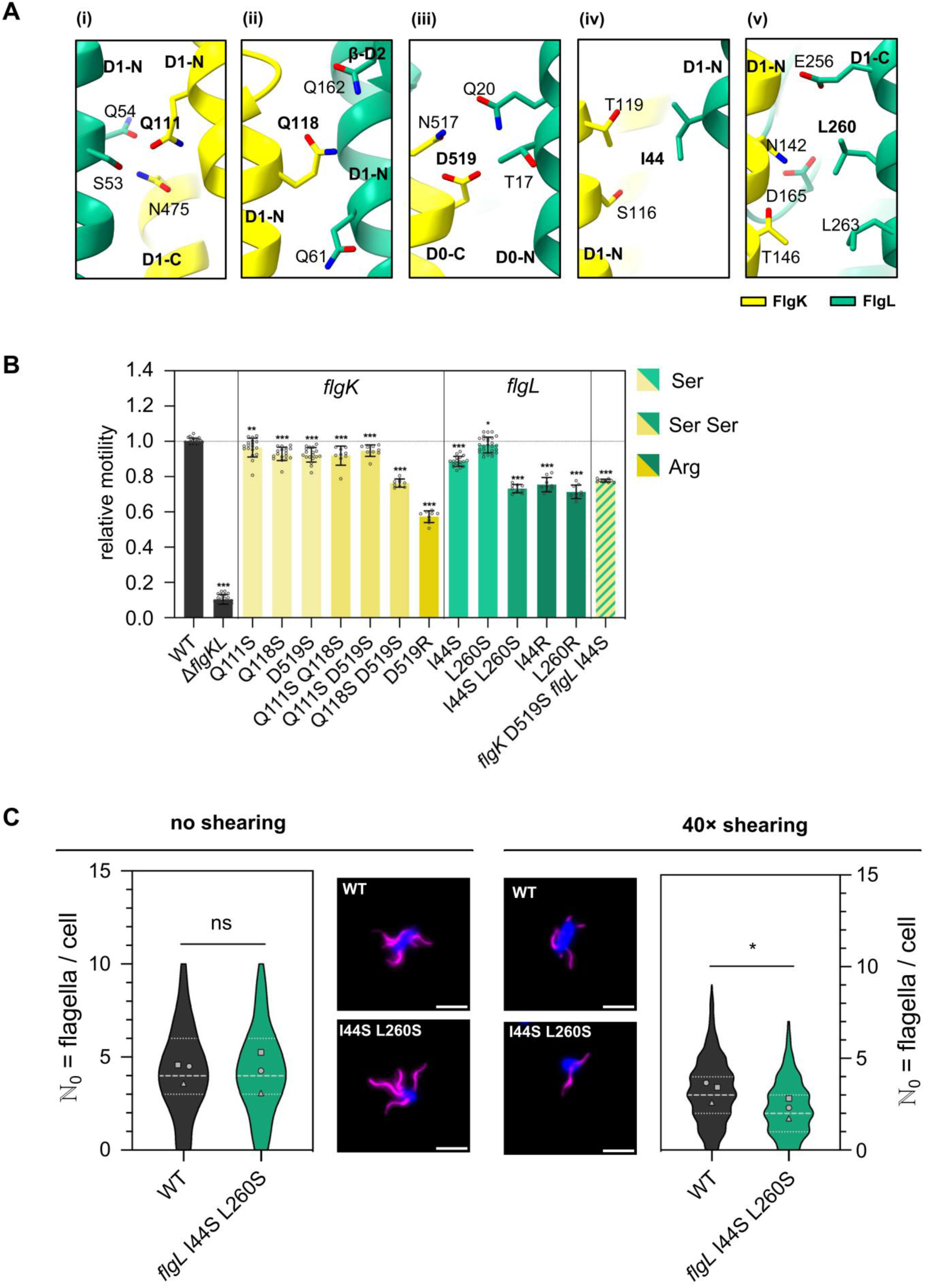
Mutations in the HFJ interface impair bacterial motility and filament stability. (A) Molecular details of interfaces between FlgK and FlgL. Residues Q111 (i), Q118 (ii), D519 (iii) for FlgK and I44 (iv), L260 (v) for FlgL are found to interact with their neighboring residues in PISA. Mutated residues and the domains they are located in, are highlighted in bold black. (B) Relative motility of *S. enterica* FlgK and FlgL point mutants analyzed using soft-agar motility plates, quantified after 4-5 h incubation at 37 °C. Diameters of the motility halos were measured using Fiji and normalized to the WT. Bar graphs represent the mean of at least 3 biological replicates with standard deviation error bars. Replicates shown as individual data points. Ser, single substitution to serine; Ser Ser, double substitution to serine; Arg, single substitution to arginine. (C) Quantification of the number of flagella per cell in the WT and FlgL I44S L260S mutant including representative fluorescence microscopy images (middle) without shearing (left) and after 40 × shearing (right) using the pulsed *flhDC* induction setup. Filaments (FliC T237C) were labeled with Dylight555 Maleimide (magenta) and DNA counterstained with DAPI. Scale bar: 2 µm. The number of flagella per cell (ℕ0) was determined for n > 150 individual bacteria per strain and condition for 3 biological replicates. Violin plots represent the distribution of the data of all replicates including the median (dashed line) and quartiles (dotted lines). Data points represent the means of each biological replicate. Statistical annotations were calculated with a two-tailed Student’s t-test, on the means of each biological replicate (*, p < 0.05; **, p < 0.01; ***, p < 0.001; ns, non-significant). See also Figure S4E and S6B.

Substitution of these FlgK residues to serine has only a mild effect on motility, with 4- 8% reduction compared to the WT (Q111S, Q118S, and D519S). When double mutations are introduced (Q111S Q118S, Q111S D519S, and Q118S D519S), only Q118S D519S shows a stronger reduction compared to the respective single serine mutants. The substitution of D519 to arginine led to the most significant phenotype, with motility reduced to 57% compared to the WT. Since the interaction of FlgK with its chaperone FlgN is proposed to occur in the C-terminal region^29^, we sought to confirm that the observed motility defect is not due to a defect in secretion (Figure S6A). Both FlgK D519S and FlgK D519R are secreted in similar amounts like WT FlgK, suggesting that the observed motility defects of the mutants did not result from impaired secretion of mutated FlgK.

For FlgL, the single serine mutants I44S and L260S show mild reductions of motility, to 88% and 97% of WT motility, respectively (Figure 5B). However, in the FlgL double mutant (I44S L260S), motility is reduced to 73% compared to the WT. Similarly, substituting the selected residues of FlgL to arginine (I44R, L260R) resulted in greater reductions to 75% and 71% WT motility, respectively. Combining the serine mutations FlgK D519S and FlgL I44S reduced the motility to 77%. In conclusion, our results indicate that the integrity of the FlgKL interface is crucial for maintaining motility in *S. enterica*. Mutations to larger, charged residues or double mutations of highly conserved residues within this interface led to significant reductions in motility, supporting our hypothesis that these interactions are vital for the stability of the HFJ. To investigate whether the mutations in FlgKL result in less stable HFJs, we performed a filament shearing assay (Figure 5C, Figure S6B) employing our pulsed induction setup (Figure S4E) to synchronize flagella synthesis. After labeling the filaments with a fluorophore-coupled maleimide dye, the cell suspension was passed through a narrow needle to break off flagellar filaments by mechanical shearing forces. Subsequently, the flagellation was analyzed using fluorescence microscopy. Without shearing, the FlgL I44S L260S mutant displays the same median number of filaments per cell as the WT (four flagella per cell). However, after 40× shearing, the WT displays a median of three flagella per cell, whereas the FlgL double mutant displays a significantly decreased median of only two flagella per cell. Additionally, the percentage of non-flagellated cells in the FlgL I44S L260S mutant doubles from 5 to 10% after shearing, while the percentage of non-flagellated cells remains approximately the same in the WT (6% vs. 5%). Furthermore, the maximum number of flagella per cell after shearing is nine in the WT and only seven in the FlgL mutant. In conclusion, these results support our hypothesis that FlgKL interface mutations lead to a less stable HFJ, as evidenced by the increased susceptibility to filament breakage under mechanical stress. This instability likely contributes to the observed motility defects, underscoring the importance of FlgKL interactions for the structural integrity and proper function of the flagellar apparatus.

### Structure of the cap complex bound to the HFJ

The cap complex was shown to assemble on the HFJ^30, 31^, which acts as a priming step to initiate filament assembly. However, the molecular details of the interaction between the HFJ and the cap at the initial stage of flagellin incorporation remain unclear. To investigate this, we aimed to determine the structures of the cap complex bound to the HFJ in the absence of the filament.

Specifically, we exploited a *C. jejuni* minicell strain with deletions in *flhG* leading to hyperflagellated cell poles, and of the flagellin genes *flaA* and *flaB*^32^. In these minicells, flagella consist of only the basal body, the hook, the HFJ, and the FliD cap (Figure 6A). We manually picked the hook tips to generate the template for the following automatic picking, leading to 79,106 particles. Particles were then subjected to several rounds of 2D classification (Figure 6B). This allowed us to produce a map of the FliD cap of *C. jejuni* assembled on the HFJ to an overall resolution of 6.5 Å with 15,077 particles (Table S1), and the atomic model of the cap-HFJ complex was built (Figure 6D).

**Figure 6.**
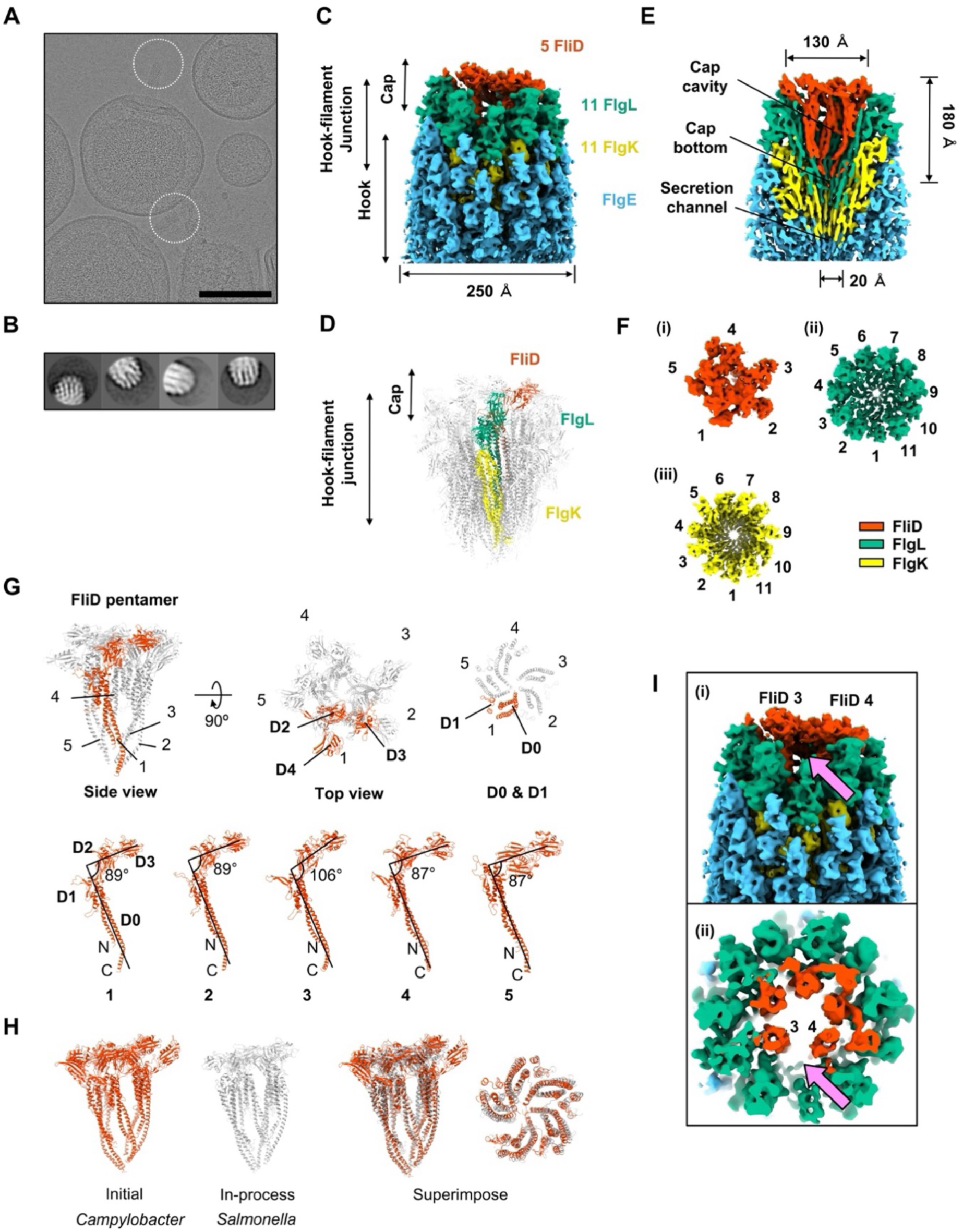
Structure of the cap complex bound to the HFJ. (A) *C. jejuni* minicells in cryo-EM showing flagellar tips to be collected. Scale bar: 200 nm. (B) 2D classification of *C. jejuni* flagellar hook tips. (C) Density map of the *C. jejuni* tip complex that contains the hook, the hook-filament junction and the cap. Components are labeled respectively, and their stoichiometry indicated. (D) Atomic model of *C. jejuni* cap-HFJ complex (PDB: 9GSX). The protofilament of FlgK-FlgL and the adjacent FliD subunit was highlighted. (E) Cross-section view of the density map of (C) with labels of dimension of FliD cap. (F) Map segment and corresponding stoichiometry of each section in the hook-junction-cap complex. (i) FliD pentamer, (ii) FlgL layer, (iii) FlgK layer. (G) Overall atomic structure of FliD cap of *C. jejuni* and structural polymorphism among FliD. (H) Superposition of the *C. jejuni* cap structure and the *S. enterica* cap structure as side view and as focus view on D0 and D1 domains. (I) Side view and cross section of the density map of *C. jejuni* tip. A gap is found between FliD 3 and FliD 4 that is primed for the first FliC to be incorporated, indicated by pink arrows. See also Movie S6.

In this structure, the cap complex also consists of a FliD pentamer with an overall width of ∼ 130 Å and a length of ∼ 180 Å - dimensions that are comparable to the cap structure bound to the filament, as described above (Figure 2D-E, **Error! Reference source not found.**E). Similar to our previous observations, the HFJ of *C. jejuni* exhibits a two-layer structure, with the proximal layer comprising 11 FlgK subunits and the distal layer comprising 11 FlgL subunits (Figure 6E, 6F). As in the structure of the filament-bound cap complex, the D0 and D1 domains of FliD shape the cap cavity, the D2 and D3 domains form the flat plane, and the D3 domain of one FliD is stacked on top of the D2 domain of another FliD in CCW direction. Notably, the *C. jejuni* FliD orthologue includes an additional domain, D4, not present in *S. enterica*, positioned to fill the gap between two FlgL subunits (Figure 6G). D4 domains are structurally distant from the cap cavity, suggesting that they are not involved in flagellin incorporation. Most FliD monomers adopt an angle of 87-89° between the D0-D1 and D2-D3 axes, which is similar to angles of 81-84° measured from the filament-bound FliD (Figure 2C, Figure 6G). However, FliD 3 exhibits an increased angle of 106°.

When superimposing the two cap complex structures, their structures are highly similar, with the major helices in D0 and D1 well aligned (Figure 6H). In contrast to the filament-bound cap structure, in the HFJ-bound cap, one of the FliD monomers is positioned downward, and its D2-D3 domains are flipped, likely corresponding to a major conformational change from initiation to elongation (Movie S6). Additional conformational changes occur in other FliD molecules, including in the D2 and D3 domains that are positioned more upward, resulting in a flat D2-D3 plane. Nonetheless, we cannot exclude the possibility that some of these structural differences may correspond to species-specific distinct conformations.

Interestingly, there is a gap between FliD 3 and FliD 4 in the cap bound to the HFJ (Figure 6I), corresponding to the position of FliC incorporation in the structure of the cap bound to the filament. We propose that this gap is the first FliC incorporation site, and therefore, the first FliC corresponds to the FliC at +11 on top of FlgL 1 (Figure 6I, S5B).

## Discussion

Here, we report the complete structure of the extracellular bacterial flagellum, encompassing the hook, HFJ, filament, and cap. Importantly, we obtain structures of assembly intermediates, which allows us to infer a complete molecular timeline for the initiation and growth of the filament. In addition, we report the structure of the cap in complex with the HFJ, presenting the HFJ as a template for filament elongation. We report mutations at respective protein-protein interfaces that impair filament assembly or structural integrity. Our structures offer critical insights into the cap-mediated incorporation of flagellins into the growing filament. Based on our structural data and mutagenesis studies, we propose a model for cap-mediated filament elongation and how the incorporation mechanism of flagellin is coupled with its secretion (Figure 7).

**Figure 7.**
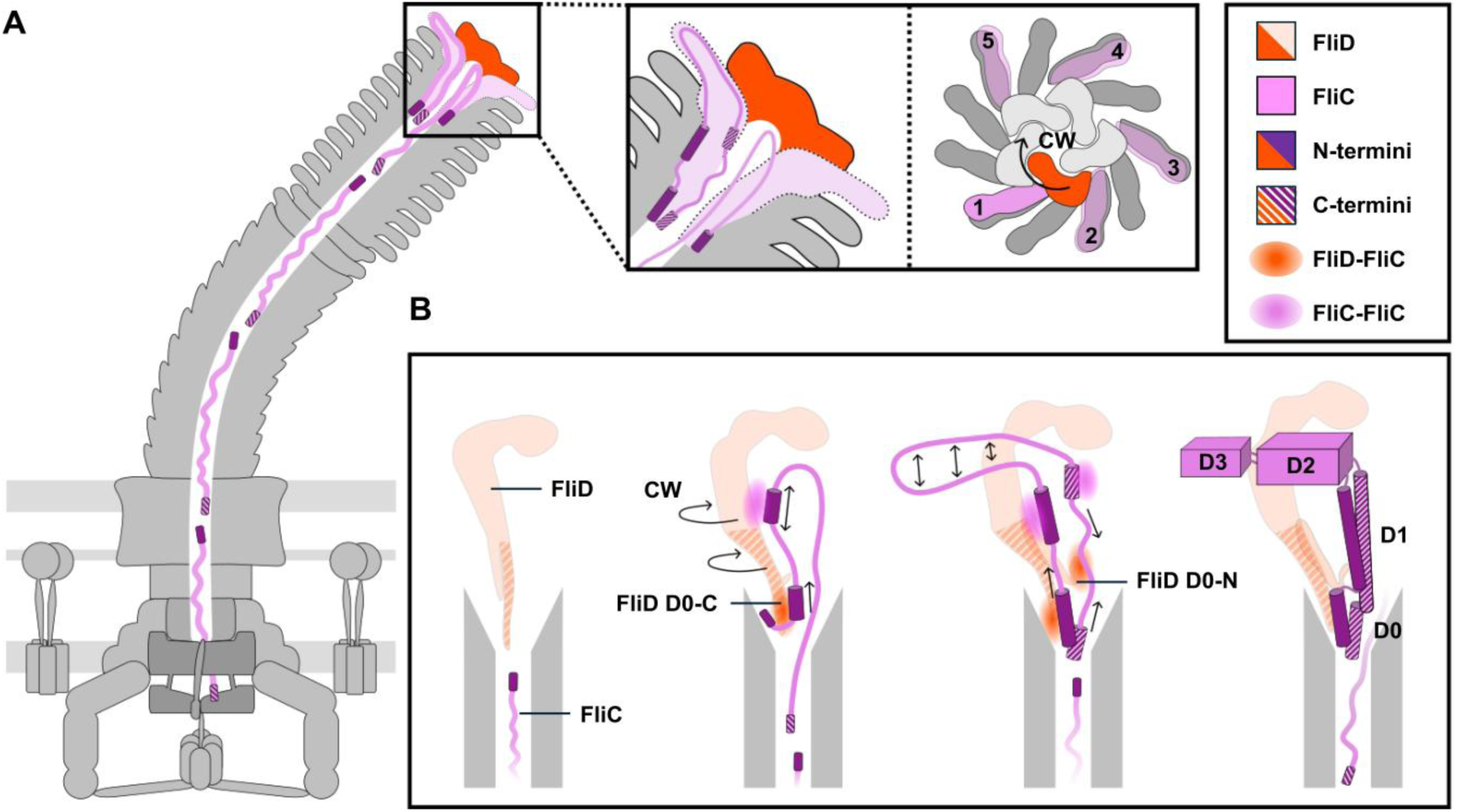
Mechanistic model of flagellin incorporation mediated by the FliD cap. (A) Flagellin incorporation at the distal end of the bacterial flagellar filament facilitated by the FliD cap. Enlarged view on the left shows a simplified cross-section of the flagellar tip, while the right side provides a top-down view. Flagellin subunits are secreted in an at least partially unfolded state, with the N- terminus leading. The incorporation of a new flagellin subunit begins before the complete maturation of the previously inserted one. The FliD cap rotates in a clockwise (CW) direction, while flagellin incorporation proceeds counterclockwise. The FliD subunit highlighted in red facilitates the insertion of the flagellin subunit labeled "1" and the prospective sites for subsequent incorporation are numbered 2-5. (B) Detailed view of the interactions involved in flagellin maturation and steps of flagellin folding. Interactions visualized as colored patches. Upon entering the FliD cap, the FliC N-terminus is captured by the D0-C domain of FliD. This interaction is followed by a 180° turn in the polypeptide chain, initiating the folding of the D0-N and D1-N domains of FliC, with stabilization provided by neighboring FliC molecules. Simultaneously, the D0, D1, and D2 domains of FliD rise, accompanied by a CW shift of the D0-D1 domains. The FliD N-terminal loop then rotates 90° upward, stabilizing FliC D0-C. Subsequent folding of FliC D1-C and D0-C occurs from both termini, with adjacent FliC subunits facilitating this process. Finally, the D2 and D3 domains are extruded from the cap cavity, completing the folding in a spontaneous manner.

### The HFJ as a functional gasket between hook and filament

Our study sheds light on the role of the HFJ, composed of 11 FlgK and 11 FlgL subunits, in the structure of the bacterial flagellum, elucidating its functional importance as a connecting platform between the flexible hook and the rigid filament.

This junction serves as a rigid linker, and based on our structural data, we suggest that it is essential for maintaining the mechanical integrity and functionality of the flagellum as a motility device. Our mutational studies confirm that the integrity and stability of the FlgKL junction plays an important role in the overall functionality of the flagellum. Altering the FlgKL interface leads to motility defects and filaments that are more prone to break (Figure 5B-C).

The FlgKL junction is crucial for bridging the highly flexible hook with the more rigid filament because without the junction proteins, FliD and FliC monomers are secreted but fail to assemble into a functional filament^30, 33^. The hook as a universal joint, which is flexible in bending yet rigid in twisting, allows for rotational movement and the transmission of torque^5, 34, 35^. In contrast, the rigid filament undergoes rapid structural changes, alternating between left-handed and right-handed helical assemblies in response to the changing direction of motor rotation^36, 37^. Earlier characterization of an *Escherichia coli* FlgL mutant suggested that the HFJ plays a crucial role in maintaining the functional shape of the filament under torsional load, thereby preventing defective motility and abnormal filament transformations^38^. Although our mutational studies did not specifically investigate this, it is consistent with our data. The separation of the hook and filament through the HFJ likely serves not only as a connecting platform but also ensures that the flexible hook can effectively transmit rotational force to the rigid filament without passing along the mechanical stress caused by its flexibility. This likely also explains why two proteins are required for the HFJ, allowing a complete isolation between the hook and the flagellins.

It is worth mentioning that the assembly mechanism of the HFJ proteins FlgKL is not currently understood. The HFJ is thought to remove the hook cap FlgD, assemble cap- less, then facilitate formation of the filament cap for filament elongation^39^. Based on our structural work, we propose that the HFJ assembles in a CW direction, as FlgK and FlgL have strong interactions along the 5-start axis but not along the 6-start axis. In addition, FlgK and FlgL interact either with copies of themselves or with adjacent proteins to which they are structurally connected. Such oriented compatibility indicates the order in which FlgK and FlgL assemble in the HFJ and that the HFJ serves as a template for filament elongation. How these proteins assemble without a cap remains to be understood.

### Mechanistic model of flagellin incorporation mediated by the FliD cap

We and others have previously proposed that the cap rotates in CW direction as a rigid pentamer with asymmetric movements of the leg domains to facilitate the incorporation of flagellins^19, 20^. In contrast, the structures reported here reveal that within the filament, the cap is asymmetric, and major structural rearrangements occur during the process of flagellin folding. Nonetheless, our data confirm that, if focusing on the entire pentamer as well as the leg domains of FliD in the incorporating state, FliD rotates in CW direction (Figure 3A, Figure 7A). Concomitantly, the incorporation of flagellin subunits occurs in CCW direction (Figure 3A, Figure 7A), consistent with earlier low resolution data^19^. Both the CW rotation of the whole FliD pentamer and individual leg domains, as well as the upward movements of the individual FliD subunits, are critical for the incorporation of flagellins into the growing filament by creating a chaperone-like environment.

Flagellins and other flagellar substrates, including FlgK, FlgL and FliD, are thought to be secreted N-terminal first via the type III secretion system due to the presence of N-terminal secretion signals^40–44^. This is fully consistent with our observation that the N-terminal D0 helix of FliC folds first. However, this secretion mechanism requires a critical 180° turn in the polypeptide chain during assembly to achieve the final tail-to-tail orientation of flagellar axial proteins^45^. Based on our data, we propose that the N-terminus of the FliC polypeptide chain interacts with the D0-C domain of one FliD subunit as soon as it emerges in the cap cavity. These interactions anchor the polypeptide chain, while the rest of the chain is secreted upwards, where the 180° turn naturally occurs (Figure 7**Error****! Reference source not found.**B). Proper folding of the D0-N and D1-N helices of FliC continues to stabilize the flagellin monomer, with our data clearly showing that the D0-N and D1-N helices are refolded subsequently after the FliC chain is anchored (Figure 3B). The D2-D3 domains of FliC are then pushed out of the cavity. Given the spatial patterns of domain appearance, we suggest that the D0-D1 domains of FliC are refolded within the FliD cavity at the incorporation site, while the D2-D3 domains are refolded outside the cap in a spontaneous manner.

Earlier studies demonstrated that the bacterial flagellar filament can assemble *in vitro* even in the absence of a cap complex; however, assembly is less efficient and requires a high concentration of purified (and potentially pre-folded) flagellin^46–48^. Importantly, many studies, including our own, have firmly established that the cap plays an essential role in ensuring filament assembly *in vivo*^18–20, 49, 50^. We conclude that the cap facilitates the incorporation of flagellin subunits by providing a chaperone-like environment that guides the proper folding and self-assembly of flagellin monomers. This efficient incorporation of flagellins mediated by the filament cap is essential, given the high metabolic cost associated with the entire process of flagella assembly. Interestingly, not only do flagellin molecules require a specialized cap structure for assembly, but so do other components such as the rod and hook, which are facilitated by the cap proteins FlgJ and FlgD, respectively^24, 51–53^. The cap structures lack sequence similarity, and their incorporation mechanisms appear to differ significantly; however, there are notable similarities. Both structures display an asymmetric organization, with lower subunits featuring high-sitting leg domains that prime the next insertion sites^24^. Hence, for the hook cap FlgD, these terminal leg domains likely play a crucial role, similar to the mechanism we propose for the function of the filament cap FliD. Therefore, both FlgD and FliD caps likely exhibit transient stabilization of assembly intermediates, ensuring proper hook or filament growth. The overall 3D structures of FlgD and FliD caps differ - for instance, terminal domains of FlgD form a helical bundle deeply embedded in the secretion channel, suggesting that the hook cap does not rotate^24^. This understanding is based on the structure of FlgD in its assembled state on top of the rod. Examining the hook cap’s active structure during hook elongation might provide valuable insights and help determine whether a rotational mechanism is conserved across flagellar cap structures. However, the molecular mechanism of the rod cap FlgJ remains even less understood and requires detailed structural characterization. Notably, FlgJ not only facilitates rod assembly but also digests the peptidoglycan layer to enable penetration through the cell wall^52, 54^. In summary, our results reported here highlight the diversity and adaptability in the strategies employed by different cap structures to facilitate flagella assembly.

### Limitations of the study

Despite providing significant insights into the structure and function of the flagellar filament cap complex, our proposed mechanism may not be entirely representative of all flagellated bacteria. The structural and functional details we report were derived from *S. enterica* and *C. jejuni*, and while FliD is conserved across many flagellated bacteria, there are proteins facilitating flagellin polymerization with analogous mechanisms, like FlaY in *Caulobacter crescentus*^55^.

## Acknowledgements

We acknowledge the support of the European Molecular Biology Organization (EMBO) through an EMBO Scientific Exchange Grant for R.E.’s research stay at the Bergeron lab. K.Q. is supported by a PhD studentship from the China Scholarship Council. M.E. acknowledges funding from the European Research Council (ERC) under the European Union’s Horizon 2020 research and innovation program (grant agreement n ° 864971) and from the Max Planck Society as Max Planck Fellow. J.R.C.B. acknowledges funding from the BBSRC (BB/R009759/2) and HFSP program (RGY0080/2021). We thank members of the Erhardt, Bergeron and Beeby labs for helpful discussions. We thank Tohru Minamino (Osaka University) for kindly providing antibodies and Christian Goosmann (Max Planck Institute for Infection Biology) for TEM grid preparations and observation of purified flagella samples for protocol optimizations. Cryo-EM grids were screened at the Imperial College London cryo-EM facility, and data was collected at the LonCEM facility; we acknowledge Paul Simpson, and Nora Cronin, respectively, for support.

## Author Contributions

J.R.C.B., M.E. and M.B. conceptualized and supervised the research project and ensured funding. R.E. prepared the purified short flagella sample of *S. enterica* and reconstructed the EM map of the FliD cap complex of *S. enterica* with the help of K.Q..

K.Q. prepared the EM grids, collected and processed the EM data. K.Q. reconstructed the EM map of the cap complex and HFJ complex of *S. enterica* and built the corresponding atomic models. R.E. and J.S. generated chromosomal *S. enterica* FliD mutants, performed and analyzed motility assays, secretion assays, and fluorescent microscopy experiments of FliD mutants. J.S. performed leakage assays. R.E. generated chromosomal *S. enterica* FlgKL mutants and performed and analyzed filament shearing assays. N.S.A. prepared samples for cryo-ET of intact *S. enterica* cells and contributed to EM grids preparation, EM data acquisition and EM data processing, with help from D.M.. T.D., E.J.C., N.G.R., J.H., E.S., M.B. designed and conducted the experiments, including generating the *C. jejuni* minicell strain, collecting and processing the cryo-EM data of *C. jejuni* minicells, and obtaining the map of the cap-HFJ complex. K.Q. built the atomic models of the cap-HFJ complex of *C. jejuni*.

R.E. and K.Q. wrote the first draft of the manuscript, prepared figures and K.Q. prepared movies. M.E. and J.R.C.B. reviewed and edited the manuscript. All authors reviewed the results and approved the final version of the manuscript.

## Declaration of interest

The authors declare no competing interests.

## STAR Methods

### Key Resource Table

| REAGENT or RESOURCE | SOURCE | IDENTIFIER |
| --- | --- | --- |
| Bacterial and Virus Strains |  |  |
| *Campylobacter jejuni* minicell stain ($\Delta flhG \Delta flaAB$) | Preprint: Drobníč et al., Cohen et al., Hendrixson et al.<sup>32, 56, 57</sup> | N/A |
| *Salmonella enterica* subsp. *enterica* serovar Typhimurium LT2 | John Roth | N/A |
| All *Salmonella enterica* mutants and control strains, see Table S2 | This study / Lab collection | N/A |
| Chemicals, Peptides, and Recombinant proteins |  |  |
| Anhydrotetracycline hydrochloride (AnTc) | Sigma-Aldrich | Cat # 13803-65-1 |
| DyLight™ 550 Maleimide | Thermo Fisher Scientific | Cat # 62290 |
| BD Difco™ Salmonellen H-Antiserum i. - Primary anti-FliC antibody | BD Difco | Cat # 11712894 |
| Secondary anti-rabbit antibody Alexa Fluor™ 488 | Invitrogen | Cat # A-11034 |
| Critical Commercial Assays |  |  |
| Q5 High-Fidelity 2× Master Mix | New England Biolabs | Cat # M0492S |
| Deposited Data |  |  |
| Cryo-ET map of the *Salmonella* flagellar tip | This study | EMDB: EMD-51555 |
| Coordinates and Cryo-EM map of the *Salmonella* cap-filament complex | This study | PDB: 9GNZ; EMDB: EMD-51486 |
| Coordinates and Cryo-EM map of the *Salmonella* hook-filament junction complex | This study | PDB: 9GO6; EMDB: EMD-51493 |
| Coordinates and Cryo-EM map of the *Campylobacter* cap-HFJ complex | This study | PDB: 9GSX; EMDB: EMD-51557 |
| Oligonucleotides |  |  |
| Primers for cloning and mutagenesis, see Table S3 | This study, Lab collection | N/A |
| Software and algorithms |  |  |
| EPU 3.4 | Thermo Fisher | https://www.thermofisher.com/uk/en/home/electron-microscopy/products/software-em-3d-vis/epu-software.html |
| CryoSPARC | Punjani et al.<sup>58</sup> | https://cryosparc.com/ |
| WarpEM v1.0.9 | Tegunov & Cramer<sup>59</sup> | http://www.warpem.com/warp/# |
| IMOD v4.11.11 | Mastronarde<sup>60</sup> | https://bio3d.colorado.edu/imod |
| Relion v4.0 | Kimanius et al., Scheres<sup>61, 62</sup> | https://relion.readthedocs.io/en/release-4.0/ |
| COOT | Emsley et al.<sup>63</sup> | https://www2.mrc-lmb.cam.ac.uk/personal/pemsley/coot/ |
| PHENIX | Liebschner et al.<sup>64</sup> | https://phenix-online.org/ |
| UCSF ChimeraX | Goddard et al.<sup>65</sup> | https://www.cgl.ucsf.edu/chimerax/ |
| ISOLDE | Croll<sup>66</sup> | https://tristanic.github.io/isolde/ |
| PyMOL | Schrodinger, Inc. | https://pymol.org/2/ |
| AlphaFold2 | Mirdita et al.<sup>67</sup> | https://colab.research.google.com/ |
| PISA | Krissinel & Henrick, Krissinel<sup>26, 27</sup> | https://www.ebi.ac.uk/pdbe/pisa/ |
| Zen 3.8 pro | Zeiss | N/A |
| Fiji 2.14.0/1.54 | Schindelin<sup>68</sup> | https://imagej.net/ |
| Image Lab | Bio-Rad | N/A |
| GraphPad Prism 10 | GraphPad Software | N/A |
| crYOLO | Wagner et al.<sup>69</sup> | https://cryolo.readthedocs.io/en/stable/index.html |
| CTFFIND-4.1 | Rohou and Grigorieff<sup>70</sup> | https://grigoriefflab.umasmed.edu/ctffind4 |
| Other |  |  |
| C-flat holey carbon films 3.5/1 200 mesh (EMS) | Electron Microscopy Sciences | https://www.emsdiasum.com |
| R0.6/1, R1.2/1.3, R2/2 holey carbon film supported copper grids, 300 mesh | Quantifoil | https://www.quantifoil.com |

## RESOURCE AVAILABILITY

### Lead Contact

Further information and requests for resources and reagents should be directed to and will be fulfilled by the Lead Contact, Julien R. C. Bergeron.

### Materials Availability

Bacterial strains generated in this study are available upon request.

### Data and Code Availability

The Cryo-ET map of *S. enterica* flagellar tip has been deposited in the EMDB database. The coordinates and EM maps including the *S. enterica* cap-filament complex, the *S. enterica* HFJ, and the *C. jejuni* cap-HFJ complex, have been deposited in the PDB and EMDB databases with the following accession code: the cryo-ET map of *S. enterica* flagellar tip, EMD-51555; the *S. enterica* cap-filament complex, PDB: 9GNZ, EMDB: EMD-51486; *S. enterica* HFJ, PDB: 9GO6, EMDB: EMD-51493; *C. jejuni* cap-HFJ complex, PDB: 9GSX, EMDB: EMD-51557.

## EXPERIMENTAL MODEL AND SUBJECT DETAILS

### *S. enterica* strains and cultivation

All *S. enterica* strains used in this study are listed in Table S3 and were derived from *S. enterica* serovar Typhimurium LT2. Bacteria were grown under constant shaking at 180 rpm in lysogeny broth (LB) at 37 °C, unless stated otherwise, and supplemented with 100 ng/mL anhydrotetracycline (AnTc) when required. Bacterial growth was determined by measuring the optical density at 600 nm using a spectrophotometer (Amersham Bioscience). For transductional crosses the general transducing *Salmonella* phage P22 HT105/1 int-201 was used^71^. Point mutations and gene deletions were introduced into the *S. enterica* genome to maintain native expression conditions using lambda-red homologous recombination^72^. The oligonucleotides used for strain construction are listed in Table S4.

### Short flagella purification

The purification of short flagella from *S. enterica* strain EM16009 (Table S3) was adapted from a previously published protocol^23–25^. Briefly, an overnight culture was inoculated with a single colony in LB medium. The next day, 500 mL LB medium was inoculated 1:100 with the overnight culture and grown for 3 h. To induce flagellin production, AnTc was added, and the cells were further inoculated for 30 min. Cells were harvested at 4,000 × g at 4 °C for 15 min. The cell pellet was carefully resuspended in 20 mL ice-cold sucrose solution (0.5 M sucrose, 0.15 M Trizma base, unaltered pH) on ice. Lysozyme and EDTA pH 4.7 were added slowly to final concentrations of 0.1 mg/mL and 2 mM, respectively, while stirring the cell suspension on ice. After 5 min of stirring on ice, the suspension was transferred to RT and slowly stirred for 1 h to allow spheroplast formation. For cell lysis, n-dodecyl β-maltoside (DDM) was added at a final concentration of 0.5%. After rapidly stirring the suspension for 10 min until it became translucent, it was slowly stirred further for 30-45 min on ice. To degrade DNA, 2 mg DNAseI and MgSO4 were added to a final concentration of 5 mM while stirring on ice. After 5 min, EDTA pH 4.7 was added at a final concentration of 5 mM. To pellet cell debris and unlysed cells, the suspension was centrifuged at 15,000 × g at 4 °C for 10 min. The supernatant was collected and centrifuged at 37,000 rpm (Beckmann Type 70 Ti rotor) for 1 h at 4 °C to pellet the flagella. The flagella were washed carefully with 30 mL buffer A (0.1 M KCl, 0.3 M sucrose, 0.05% DDM) and collected again at 37,000 rpm (Beckmann Type 70 Ti rotor). Flagella were resuspended carefully in 100 µL buffer B (10 mM Tris pH8, 5 mM EDTA pH 8, 0.01% DDM) and incubated overnight on a rolling platform at 4 °C. The next day, samples were used for cryo-EM grid preparation.

### Single-particle cryo-EM sample preparation and data acquisition of *S. enterica* cap and hook-filament junction

Short flagella resuspension (3 μl) was applied to glow-discharged holey carbon grids (Quantifoil R2/2, 300 mesh). Samples were incubated for 30 s at 4 °C and 88% humidity before being blotted by Leica EM GP1 and then rapidly plunged into liquid ethane. Grids that were blotted for 3, 4, 5, 6 s were screened on 200 kV Glacios microscope (Thermo Fisher). The grids with good ice thickness were deposited to 300 kV Krios G3i microscope with a Gatan K3 direct electron detector (Thermo Fisher). Dataset was collected using a physical pixel size of 1.078 Å at the magnification of 81,000 ×. Finally, 24,729 movies were collected at a dose rate of 16.9 e-/pix/s and exposure of 3.2 s, corresponding to a total dose of 43 e-/Å^2^. All movies were collected over 40 frames with a defocus range of −0.9 μm to -2.7 μm.

### Single-particle cryo-EM image processing and EM map reconstruction of *S. enterica* cap and hook-filament junction

For all the movies, the motion-correction and the CTF estimation were processed in CryoSPARC v4.4.^58^ using the patch motion correction and the patch CTF estimation respectively. A total of 24,066 movies were used for the following data processing.

For FliD cap, 1,896 particles were manually picked and subjected to 2D classification to generate templates for the following template-based automatically picking. 527,490 particles were recognized and extracted with a box size of 500×500 pixels. Multiple rounds of 2D classification were performed to purify the dataset in which the particles that either are not FliD cap or with low quality were excluded. A subset of 63,208 particles was selected in 2D classification. After a homogeneous refinement, local CTF refinements and a round of non-uniform refinement, an overall 3.28 Å resolution density map of the FliD cap complex was obtained. 3D variability analysis^28^ was performed to analyze discrete heterogeneity and resolve continuous flexibility. To further resolve the density of the D2-D3 of the most flexible FliD, 3D classification with 15 classes was performed attempting to discriminate different transition states of FliD cap during elongation, which resulted in 8 different states of FliD cap. A subset of 15,225 particles was selected from the best class followed by non-uniform refinement and local refinement with a mask where D2-D3 domains of flagellin are removed, an average 3.74 Å resolution density map with all FliD subunits well-resolved was obtained.

For hook-filament junction, filament tracer was performed firstly with a filament diameter of 220 Å and a separation distance of 0.25-fold diameters. A subset of flagellin filament obtained from the initial 2D classification of FliD cap was used as the template for filament tracer. 3,349,411 particles were extracted with a box size of 500×500 pixels and subjected to 2D classification. Several rounds of 2D classification were performed to remove the cap, the hook, the filament and the basal body, and a subset of 125,307 particles were selected. After a homogeneous refinement, a round of local CTF refinement and a round of non-uniform refinement, an average 2.54 Å resolution density map of the junction complex was obtained, where the density of the FlgKL junction is mixed with the density of FlgE hook and FliC filament. Next, 3D classification with 15 classes was performed to discriminate junctions at different positions relative to the density map and only two junction classes were obtained, 65,561 particles and 58,875 particles respectively. After performing non-uniform refinement for each class, an average 3.02 Å and 3.23 Å resolution density map of junction with less mixing signal of hook and filament of each class was obtained. A local refinement with a tight mask was performed based on the 3.02 Å junction map, which improved the average resolution of the junction to 2.91 Å and the occupancy of FlgK and FlgE.

### Atomic model building and refinement of *S. enterica* cap and hook-filament junction

For FliD-FliC model, the monomer structure of full-length FliD and FliC were generated using AlphaFold2^67^. First, the D0-D1 domain and D2-D3 domain of each FliD were isolated, manually fitted into the reconstructed maps as rigid bodies, and the FliD pentamer was built. Flexible fitting was then performed on the FliD pentamer using ISOLDE^66^ with secondary structure restraints gained from the AlphaFold model in UCSF ChimeraX^65^. After the flexible fitting completed, D0-D1 and D2-D3 of each FliD was connected automatically in PyMOL. FliC flagellins were then replicated and fitted into the map as rigid bodies manually. ISOLDE was used again to flexibly fit the whole complex of 5 FliD and 17 FliC into the reconstructed map. The FliD-FliC complex model was refined with real-space refinement in PHENIX^64^ with secondary structure, rotamer and Ramachandran restraints but without NCS restraints. Coot was then used to correct rotamer outliers and side-chain clashes and was also used to delete residues with no density can fit, and vice versa^63^. The final model was validated using comprehensive validation program with default settings in PHENIX. For the hook-filament-junction model, monomer structures of the full-length FlgE, FlgK, FlgL and FliC were generated using AlphaFold2^67^. All subunits were manually fitted into the reconstructed maps as rigid bodies at first, including 13 FlgE, 11 FlgK, 11 FlgL and 14 FliC. Then ISOLDE was used for flexible fitting with secondary structure restraints for the entire complex, followed by the same refinement and validation procedures and settings as we refined the FliD-FliC model using PHENIX and Coot.

### Swimming motility

Swimming motility was studied using tryptone broth-based soft agar swim plates containing 0.3% Bacto agar. Motility plates were inoculated with 2 µL of overnight culture and incubated at 37 °C for 4-5 h. Images were acquired by scanning the plates and the diameters of the swimming halos were measured using Fiji^68^. The swimming diameters of the mutant strains were normalized to those of the WT.

### Pulsed *flhDC* induction setup

To synchronize flagellar biosynthesis and, when required, prevent the negative feedback loop of FliT on FlhDC, we used strains with an AnTc inducible promoter for the master regulator FlhDC (P*tetA*-*flhDC*). Flagellum formation is regulated by multiple feedback loops, involving the chaperone FliT^73–76^. In the cytosol, FliT binds to FliD, targeting it to the export gate. Upon HBB completion and secretion of FliD, FliT is released and binds to the FlhC subunit of the flagellar master regulator, preventing expression of flagellar class 2 and subsequently class 3 genes^74–76^. Previous studies have shown that the C-terminal region of FliD binds FliT^77–79^. To minimize putative effects of the FliT feedback loop in our C-terminal FliD mutants and study flagellin polymerization, we synchronized flagella assembly by expressing *flhDC* from an inducible promoter (P*tetA-flhDC*)^80^. We reasoned that cellular levels of free FliT would be low during initial rounds of flagella assembly, as unbound FliT accumulates over time.

Briefly, we induced *flhDC* expression with a 30 min pulse of anhydrotetracycline (AnTc), followed by removal of the inducer and a 60 min incubation step (Figure S4E). This pulsed *flhDC* induction setup was used for fluorescence microscopy, flagellin leakage, filament shearing, and secretion assays (Figure S4E). Overnight cultures were grown in the absence of AnTc and were diluted 1:10 in a total volume of 10 mL at 30 °C for 1.5 h before inducing flagellar biosynthesis with the addition of AnTc. Cells were grown for 30 min in the presence of AnTc at 30 °C. Cells were centrifuged for 5 min at 2,500 × g to remove the inducer, resuspended in 10 mL fresh LB, and incubated further for 60 min at 30 °C before sample collection.

### Fluorescence microscopy

All strains investigated by fluorescence microscopy, except for EM16009 that was used for short flagella purification, were locked in the expression of *fliC* (Δ*hin*- 5717::FRT) and contained a cysteine mutation in FliC (*fliC*6500 T237C), which allowed labeling of the flagellin subunit with a fluorophore-coupled maleimide dye^81^. After culturing the cells using the pulsed *flhDC* induction setup, 500 µL of cell suspension was collected. Cells were centrifuged for 5 min at 2,500 × g and resuspended in 500 µL of 1 × PBS. Fluorophore-coupled maleimide dye (Alexa Fluor 488, Invitrogen) was added at a final concentration of 10 μM, and cells were further incubated for 30 min at 30 °C. Following the removal of unbound dye by slow-speed centrifugation for 5 min at 2,500 × g, cells were resuspended in 500 μl 1 × PBS and applied to a homemade flow cell. Flow cell preparation was performed as previously described^82, 83^. Briefly, coverslips were incubated with 0.1% poly-L-lysine (PLL) for 10 min, air-dried for 10 min, and subsequently fixed to an objective slide via two layers of pre-heated parafilm to create a chamber. The side of the coverslip incubated with PLL faced the objective slide. Cells were allowed to adhere to the coverslip for 3 min at room temperature in the dark and were subsequently fixed with 2% (v/v) formaldehyde and 0.2% (v/v) glutaraldehyde for 10 min. Cells were washed twice with 1 × PBS and finally mounted with Fluoroshield™ + DAPI solution (Sigma-Aldrich). Images were acquired by an inverted epifluorescence microscope (Zeiss AxioObserver.Z1) at 100 × magnification with a Prime BSI Scientific CMOS (sCMOS) camera, a Pln Apo 100×/1.4 Oil Ph3 objective and a LED Colibri 7 light source (Zeiss). Images were taken using the Zen 3.8 pro software with Z-stack every 0.5 µm with a range of 3 µm (7 slices). Images were analyzed using Fiji^68^ and the NeuronJ plugin^84^.

To test the optimal AnTc induction time for flagellin production and short flagella purification (Figure S1D), an overnight culture of EM16009 was diluted 1:100 and grown for 3 h before induction of FliC production with AnTc for 30, 45, 60 or 120 min. At each time point, 500 µL samples were collected, loaded onto a flow-cell and fixed as described above. Cells were washed twice with 1 × PBS and blocked with 10% BSA for 10 min. Primary α-FliC antibody (BD Difco™ Salmonellen H-Antiserum i., catalog number 11712894, 1:1K in 2% BSA) was added for 1 h. Cells were washed twice with 1 × PBS, and subsequently blocked with 10% BSA for 10 min. Secondary antibody (anti-rabbit-Alexa488, invitrogen, catalog number A-11034, 1:1K in 1 × PBS) for 30 min. Cells were washed twice with 1 × PBS and mounted with Fluoroshield™ + DAPI solution (Sigma-Aldrich). Fluorescence microscopy was carried out as described above.

### Flagellin leakage assay

After culturing the cells using the pulsed *flhDC* induction setup, a 3 mL aliquot was taken, and cells were harvested by centrifugation at 13,000 × g for 5 min. The samples were further treated to separate monomeric FliC from cytoplasmic, cell-associated, and cell-detached FliC (Figure 4E, S4E). 2 mL of the supernatant were transferred to a fresh tube, the remaining supernatant was removed, and the cell pellet was resuspended in 1.5 mL 1 × PBS. The cell pellets were incubated at 65 °C for 5 min to depolymerize the flagellar filament, and the samples were subsequently centrifuged to obtain cell pellets and supernatants containing cytoplasmic flagellin molecules and depolymerized flagellin monomers, respectively. The supernatants were ultracentrifuged at 85,000 × g for 1 h at 4 °C, and the pellets containing flagellar filaments detached from the cell bodies and supernatants containing flagellin monomers leaked into the culture medium were collected separately. Proteins from different fractions were precipitated using 10% trichloroacetic acid (TCA) and separated by SDS-PAGE. FliC protein levels in the different samples were determined by immunoblotting using primary α-FliC (Difco, catalog number 228241 *Salmonella* H Antiserum I, 1:5,000 in 1 × TBS-T) and secondary α-rabbit antibodies (Bio-Rad Immun-Star Goat Anti-Rabbit (GAR)-HRP Conjugate, catalog number 170-5046, 1:20,000 in 1 × TBS-T). Relative FliC protein levels were normalized to the housekeeping protein DnaK, which was detected using α-DnaK (abcam) antibody and secondary α-mouse antibodies conjugated to horseradish peroxidase (Bio-Rad Immun-Star Goat Anti-Mouse (GAM)-HRP Conjugate, catalog number 170-5047, 1:20,000 in 1 × TBS-T) antibodies, using the Image Lab software (Bio-Rad). The percentage of flagellin monomers secreted into the culture supernatant was calculated by dividing the amount of secreted flagellin monomers by the total flagellin amount, comprising secreted and cellular as well as detached and attached flagellin molecules.

### Filament shearing assay

All strains investigated in the shearing assay were locked in the expression of *fliC* (Δ*hin*-5717::FRT) and contained a cysteine mutation in FliC (*fliC*6500 T237C), which allowed labeling of the flagellin subunit with a fluorophore-coupled maleimide dye^81^. After culturing the cells using the pulsed *flhDC* induction setup, 1000 µL of the cell suspension was collected. Cells were centrifuged for 5 min at 2,500 × g and resuspended in 500 µL of 1 × PBS. Fluorophore-coupled maleimide dye (Alexa Fluor 488, Invitrogen) was added at a final concentration of 10 μM, and cells were further incubated for 30 min at 30 °C. Following the removal of unbound dye by slow-speed centrifugation for 5 min at 2,500 × g, the cells were resuspended in 1000 μL 1 × PBS. Samples were split into 2 × 500 µL aliquots. One sample was left untreated, while the flagella of the other sample were sheared by passing the cell suspension 40 × back and forth through a 27G, 0.4x12mm BL/LB needle using a 1 mL syringe. For the first replicate (Figure S6B), the samples were split into 4 × 500 µL aliquots. Again, one sample was left untreated, while the flagella of the other samples were sheared by passing the cell suspension 20 ×, 40 ×, or 60 × back and forth through a 27G, 0.4x12mm BL/LB needle using a 1 mL syringe. To separate the cells from the sheared filaments, the samples were centrifuged at 2,500 × g, and the supernatant containing most of the sheared filaments was discarded. Cells were resuspended in 500 µL of 1 × PBS and applied to a homemade flow cell as described above. Images were acquired by an inverted epifluorescence microscope (Zeiss AxioObserver.Z1) at 100 × magnification with a Prime BSI Scientific CMOS (sCMOS) camera, a Pln Apo 100×/1.4 Oil Ph3 objective and a LED Colibri 7 light source (Zeiss). Images were taken using the Zen 3.8 pro software with Z-stack every 0.5 µm with a range of 3 µm (seven slices). Images were analyzed using Fiji^68^.

### Protein secretion assay

After culturing the cells using the pulsed *flhDC* induction setup, a 1.9 mL aliquot was taken, and cells were harvested by centrifugation at 13,000 × g for 5 min. 1 mL of the supernatant was transferred into a fresh tube, the remaining supernatant was discarded, and the cell pellets were resuspended in 1 mL of double-distilled water. Proteins were precipitated from the supernatant and pellet fractions using 10% (v/v) TCA, followed by a 30 min incubation step on ice and centrifugation at 20,000 × g for 30 min. Protein pellets were washed with ice-cold acetone and air-dried. Samples were adjusted to 20 OD units/µL, and 200 OD units were analyzed under denaturing conditions using SDS-PAGE. Immunoblotting was performed using primary α-FliC (Difco, catalog number 228241 *Salmonella* H Antiserum I, 1:5,000 in 1 × TBS-T), α- FliD (Gift from T. Minamino, 1:10,000 in 1 × TBS-T) or α-FlgK (Gift from T. Minamino, 1:10,000 in 1 × TBS-T) antibodies. Proteins were detected using secondary α-rabbit antibodies conjugated to horseradish peroxidase (Bio-Rad Immun-Star Goat Anti-Rabbit (GAR)-HRP Conjugate, catalog number 170-5046, 1:20,000 in 1 × TBS-T). The relative amounts of secreted and cellular proteins were determined by normalization to the housekeeping protein DnaK using the Image Lab software (Bio-Rad). DnaK was detected using a primary α-DnaK (abcam) antibody and secondary α-mouse antibodies conjugated to horseradish peroxidase (Bio-Rad Immun-Star Goat Anti-Mouse (GAM)-HRP Conjugate, catalog number 170-5047, 1:20,000 in 1 × TBS- T).

### Statistical analyses

Statistical analyses were performed using GraphPad Prism 10 (GraphPad Software, Inc., San Diego, CA), and values of P < 0.05 were considered statistically significant.

### Cryo-ET sample preparation of *S. enterica* FliD cap

An overnight culture of strain EM8327 (Table S3) in LB was diluted 1:100 in 10 mL of LB and incubated for 1.5 h. AnTc was added to induce flagellar biosynthesis and the cells were further incubated for 45 min. Cells were centrifuged at 2,550 × g for 2 min and resuspended in remaining LB. A 5% BSA solution was used to resuspend the pellet and left at 4 °C for 30 min, after which the sample was centrifuged at 6,000 × g for 10 min, and the pellet was resuspended in 200 µL of 50 mM HEPES, 100 mM NaCl pH 7 buffer. This was followed by an additional centrifugation step under the same conditions and resuspension in 35 µL buffer. Colloidal gold solution (1 mL, 5 nM) was centrifuged for 10 min at 6,000 × g. The pipette tip was cut off to allow larger cells to pass through onto the grid undamaged, and 10 µL of sample was mixed with 10 µL of colloidal gold solution prior to loading 5 µL of the mixture onto C-flat holey carbon films 3.5/1 200 mesh (EMS) using a Leica EM GP plunge freezer. Cryo-EM grids were negative glow discharged for 30 s and the double blotting strategy was used, where 5 µL of sample was loaded onto the grid for 2 min at 80% humidity and 4 °C chamber conditions, back-blotted, loaded once more for 2 min, back-blotted and loaded for 1 min, and back-blotted for 6.5 seconds before plunging into liquid ethane to freeze the grid.

### Cryo-ET data collection and processing of *S. enterica* FliD cap

Cryo-EM tomography data were collected using a Titan Krios TEM (Thermo Fisher) operated at 300 kV and equipped with a Falcon IV camera. 68 tomograms were collected using the EPU software (Thermo Fisher) in linear mode, with a pixel size of 3.6 Å pix^−1^, with a total dose of 86 e^−^ Å^−2^ spread across 35 tilts with 10 fractions each in 3 degree increments with a range of −51 to +51 degrees. The defocus range used for data collection was approximately −2 μm to −8 μm.

Pre-processing was performed using the WarpEM software package v1.0.9, and aligned using IMOD v4.11.11^59, 60^. Particle picking was performed manually on binned and deconvoluted tomograms using the 3Dmod software. Coordinates were transferred to a (.star) file and sub-tomogram reconstruction was performed in WarpEM and imported to Relion v4.0 with 252 sub-tomograms^59, 61, 62^.

The flagellum filament model (EMD-9896) was chosen as a starting model, low-pass filtered to 60 Å, and subjected to 3D refinement in Relion v4.0, with C1 symmetry, generating *de novo* density for the tip^62^. Further 3D classification and CTF refinement in the WarpEM software did not further increase the map resolution, presumably because of the limited number of sub-tomograms.

### *C. jejuni* strain construction and cultivation

A *C. jejuni* minicell (Δ*flhG* Δ*flaAB*) strain was constructed as described previously^56, 57^. Briefly, *aphA*-*rpsL*^WT^ cassettes flanked by ∼500 bp overhangs with homology to the targeted chromosomal loci and ecoRI sites at the 5’ and 3’ termini were synthesised by “splicing by overlap extension” PCR (SOE PCR). Linear DNA fragments were methylated at their ecoRI sites with ecoRI methyltransferase (New England Biolabs) and transformed into *C. jejuni* using the biphasic method^85^. Transformants were selected for on MH agar supplemented with 50 µg/mL kanamycin. Replacement of the *aphA*-*rpsL*^WT^ with the desired mutation was achieved using the same method, but with transformants being selected for on MH agar supplemented with 2 mg/mL streptomycin sulfate. Kanamycin-sensitive, streptomycin-resistant transformants were single-colony purified and checked by Sanger sequencing (Source Biosciences UK). For the minicell background, in-frame deletion of *flhG* leaves the first and last 20 codons intact, while the Δ*flaAB* allele spans from 20 base pairs upstream of the *flaA* translational start site to codon 548 of *flaB*.

### Cryo-EM sample preparation of *C. jejuni* minicell

*C. jejuni* Δ*flhG* Δ*flaAB* cells were grown on MH plates and resuspended in phosphate-buffered saline (PBS buffer, 137 mM NaCl, 2.7 mM KCl, 10 mM Na2HPO4, 1.8 mM KH2PO4, pH 7.4). Cells were spun at 1,500 × g for 20 min to pellet whole cells. The minicell-enriched supernatant was removed and spun in a tabletop microcentrifuge at 15,000 × g for 5 min to pellet the minicells. The pellet was then resuspended to a theoretical OD600 of ∼15.

Minicells were vitrified on QUANTIFOIL® R0.6/1 or R1.2/1.3 holey carbon grids (Quantifoil Micro Tools) using a Vitrobot Mark IV (Thermo Fisher Scientific).

### Single-particle cryo-EM data collection and processing of *C. jejuni* cap-hook-filament junction complex

Micrographs of the minicell sample were collected using 300 keV Thermo Fisher Scientific Titan Krios TEMs using EPU acquisition software with a K2 direct electron detector equipped with a GIF energy filter (Gatan), using a slit width of 20 eV. Due to our large particle size relative to that of the holes, we collected one shot per hole. Gain correction was performed on-the-fly.

Particle coordinates were found using a crYOLO^69^ model derived from manually picked particles from 2718 micrographs and a box size of 180 nm. Processing was done in RELION 4.0^61^. Contrast transfer function (CTF) correction was performed using CTFFIND-4.1^70^. A total of 79,106 particles was extracted from 42,988 micrographs (2.2 Å/pix). After several rounds of 2D classification and a 3D classification with a 500 Å-diameter mask, particles were re-centered at the tip of the axial structure and re-extracted. Additional rounds of 2D classifications and a 3D classification with a 400 Å-diameter mask with no symmetry imposed and post- processing to yield the final map.

## Notes

### Competing Interest Statement

The authors have declared no competing interest.

