## Supplementary figures and tables for "Structure of the complete extracellular bacterial flagellum reveals mechanism for flagellin incorporation"

### Supplemental Figures

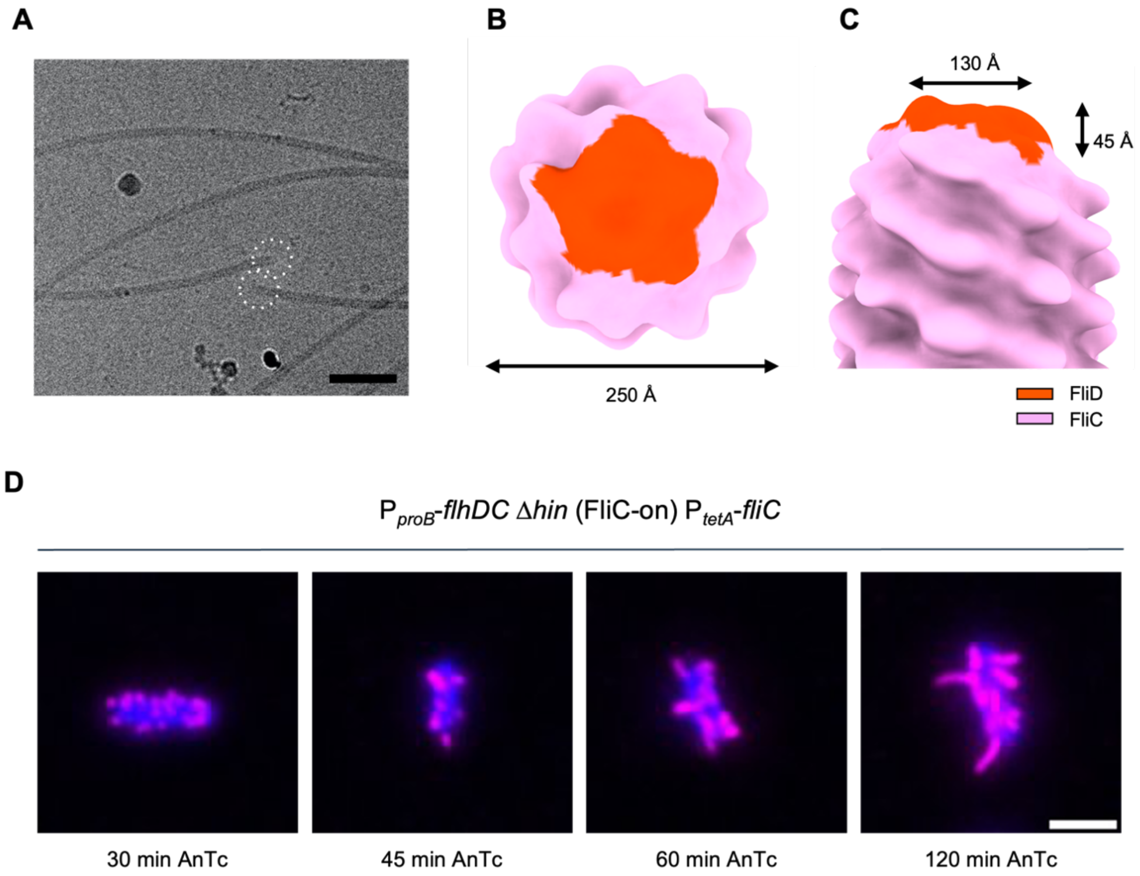


**Figure S1. Cryo-ET of filament tips and test of different flagellin induction times for short flagella purification, related to Figure 1.**
(A) *S. enterica* flagella in cryo-EM. Scale bar: 100 nm.
(B) Top view of flagellar tip with the diameter of filament.
(C) Side view of flagellar tip with the dimensions of the FliD cap.
(D) Representative fluorescence microscopy images of *S. enterica* EM16009 that was used for short flagella purification. Filaments (FliC) were immunostained with anti-FliC primary antibody and anti-rabbit coupled to GFP after inducing flagellin production with anhydrotetracycline (AnTc) for 30, 45, 60 or 120 min. DNA counterstained with DAPI. Color of the GFP channel changed to magenta. Scale bar: 2 μm.

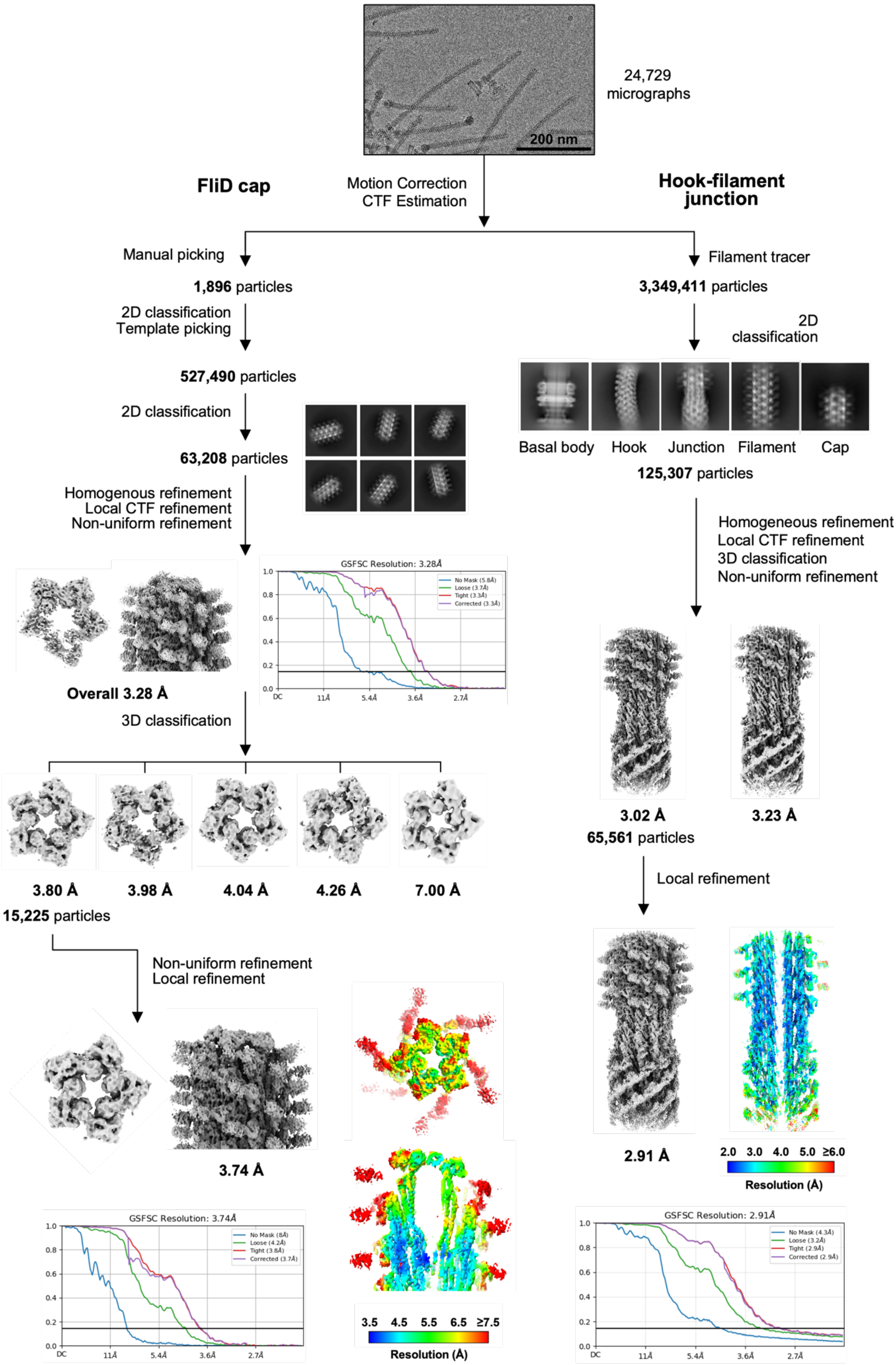


*(legend on next page)*

**Figure S2. Cryo-EM data processing pipeline, related to Figure 1.**
Flowchart of data collection and processing pipeline in cryoSPARC that resulted in the final *S. enterica* HFJ and cap complex cryo-EM structure. FliD cap: 1,896 particles were manually picked to generate a template. Template-based picking resulted in 527,490 particles from 24,729 micrographs. After 2D classification, 63,208 particles were used to generate and refine an initial model, obtaining a 3.28 Å resolution map. 3D classification was performed to solve heterogeneity in the sample. The largest class containing 15,225 particles was further refined, obtaining a 3.74 Å resolution map and cryo-EM density maps coloured by local resolution (in Å) of the top-view and cross-section of the FliD cap in complex with the filament estimated in cryoSPARC shown. Flowchart includes Gold-Standard Fourier Shell Correlations of the initial map with an estimated global resolution of 3.28 Å at FSC = 0.143 and of the final map with an estimated resolution of 3.74 Å at FSC = 0.143. Hook-Filament Junction: The use of Filament Tracer led to an initial set of 3,349,411 particles from the same 24,729 micrographs. After 2D classification, 125,307 particles were used to generate an initial map. After 3D classification and refinements of the largest class containing 65,561 particles, the map reached a resolution of 3.02 Å. Local refinement further improved resolution to 2.91 Å and cryo-EM density map coloured by local resolution (in Å) of the cross-section of the FlgKL junction in complex with the hook and filament estimated in cryoSPARC shown. Flowchart includes Gold-Standard Fourier Shell Correlation of the final map with an estimated global resolution of 2.91 Å at FSC = 0.143.

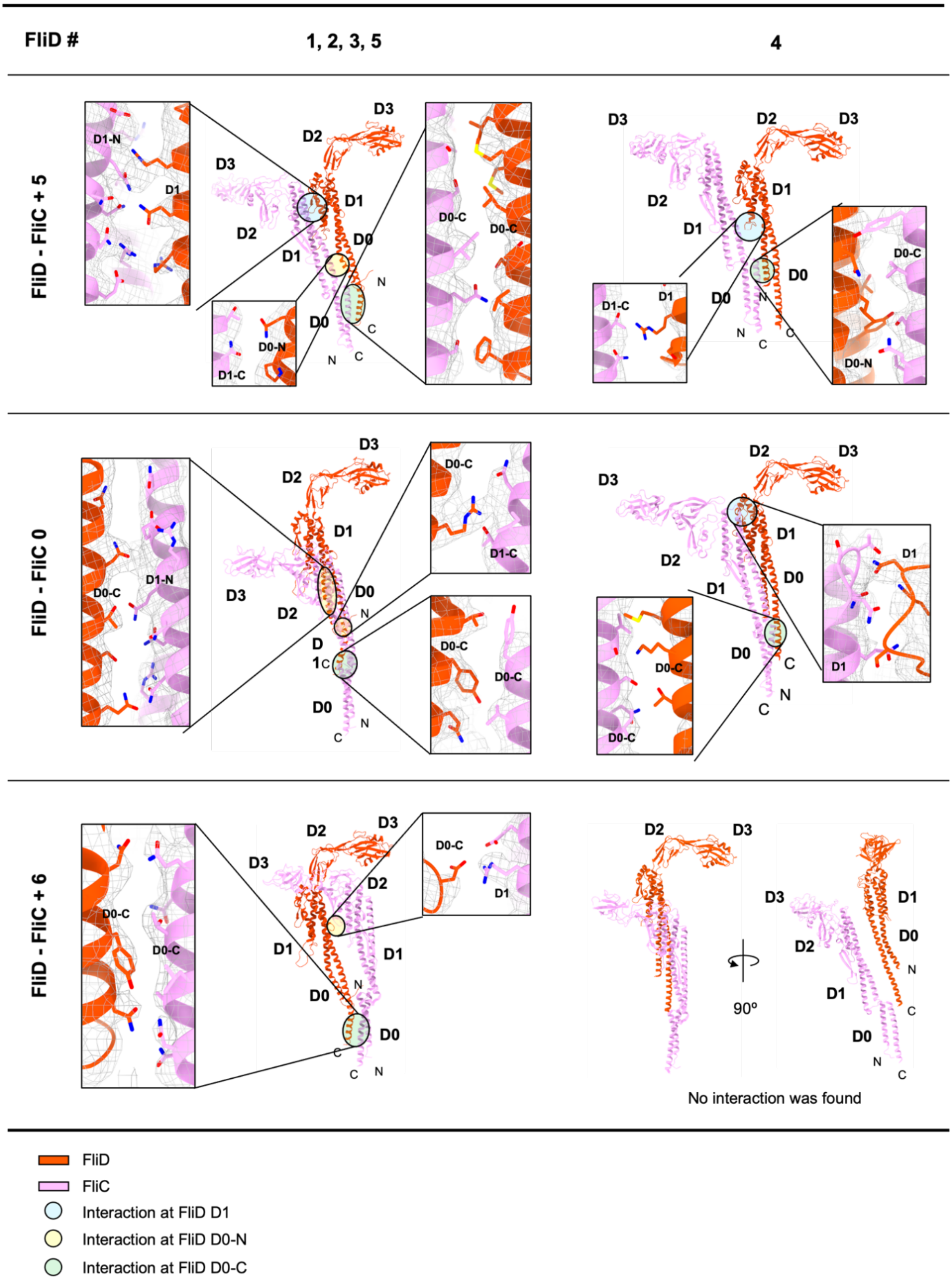


**Figure S3. Details of the FliC-FliD interactions, related to Figures 2 and 3.**
The interface between each FliD and FliC at position +5, 0, +6 are displayed and residues that are potentially interacting are displayed. FliD 1, 2, 3, 5 are grouped as they adopt similar interactions with adjacent FliC.

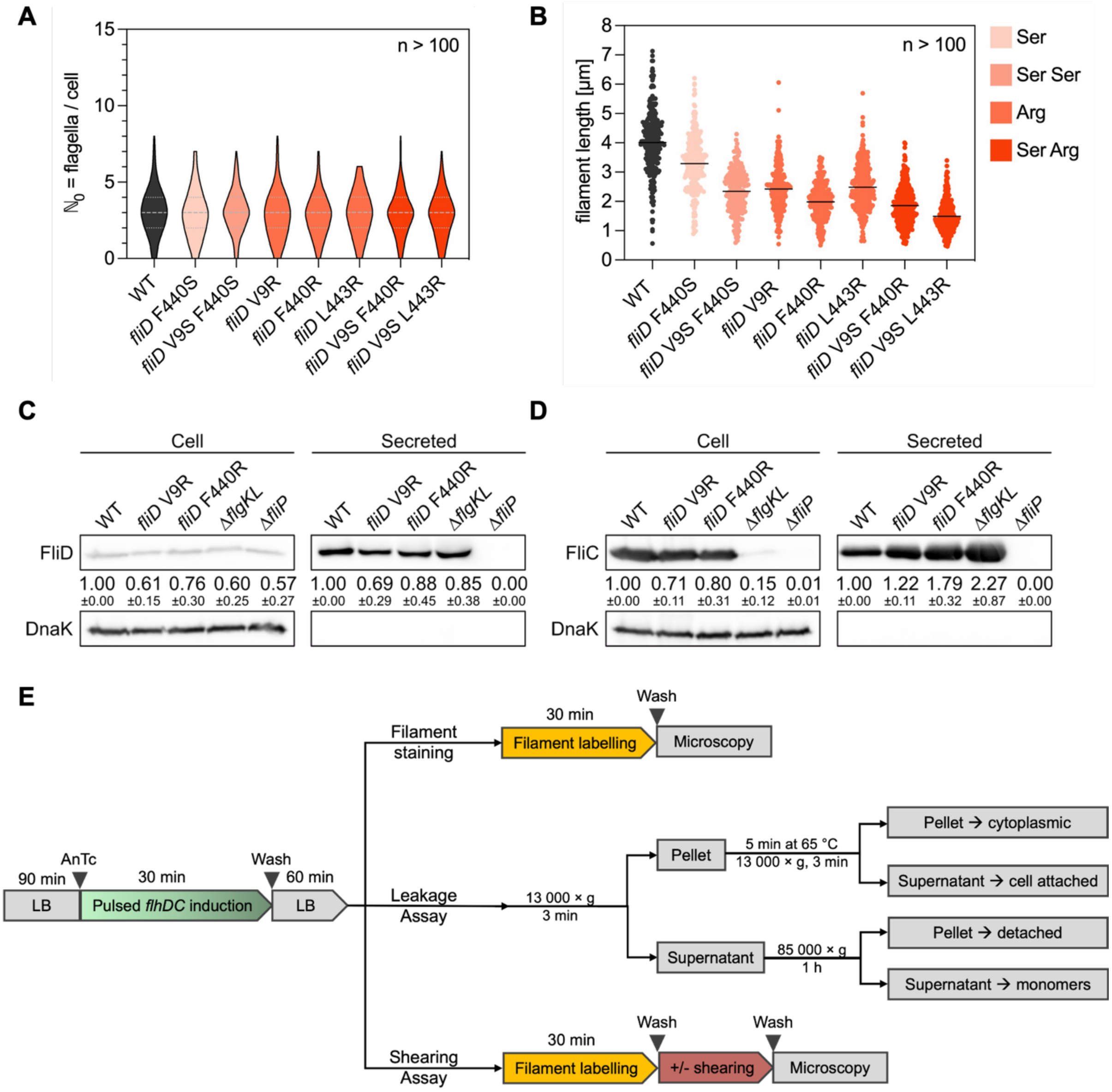


**Figure S4. Comparative analysis of filament characteristics and protein secretion in wild-type and FliD mutant strains, related to Figure 4-5 and STAR Methods.**

(A) Quantification of the number of flagella per cell in the wild-type and FliD mutants using the pulsed *flhDC* induction setup. The number of flagella per cell ($\mathbb{N}_0$) was determined for n > 100 individual bacteria per strain for one biological replicate. Violin plots represent the distribution of the data including the median (dashed line) and quartiles (dotted lines).

(B) Quantification of the filament length in µm in the wild-type and FliD mutants using the pulsed *flhDC* induction setup. The filament length was determined for n > 100 individual filaments per strain for one biological replicate. All data points shown, black lines indicate the means. Ser, single substitution to serine; Ser Ser, double substitution to serine; Arg, single substitution to arginine; Ser Arg, double substitution to serine and arginine.

(C + D) Immunoblotting of the cellular and secreted fractions using the pulsed *flhDC* induction setup of wild-type, the FliD mutants (V9R and F440R), a Δ*flgKL* strain and a secretion-deficient Δ*fliP* mutant using (C) anti-FliC or (D) anti-FliD antibodies. Relative secreted FliC or FliD levels report mean ± standard deviation, n = 3. DnaK serves as a loading and lysis control and was used to normalize the protein levels.

(E) Pulsed *flhDC* induction setup used to analyze filament number and filament length with fluorescence microscopy, and perform secretion, leakage and shearing assays. For secretion assays, the first Pellet and Supernatant samples of the Leakage assay were directly used for TCA precipitation and SDS-PAGE analysis, without the additional steps.

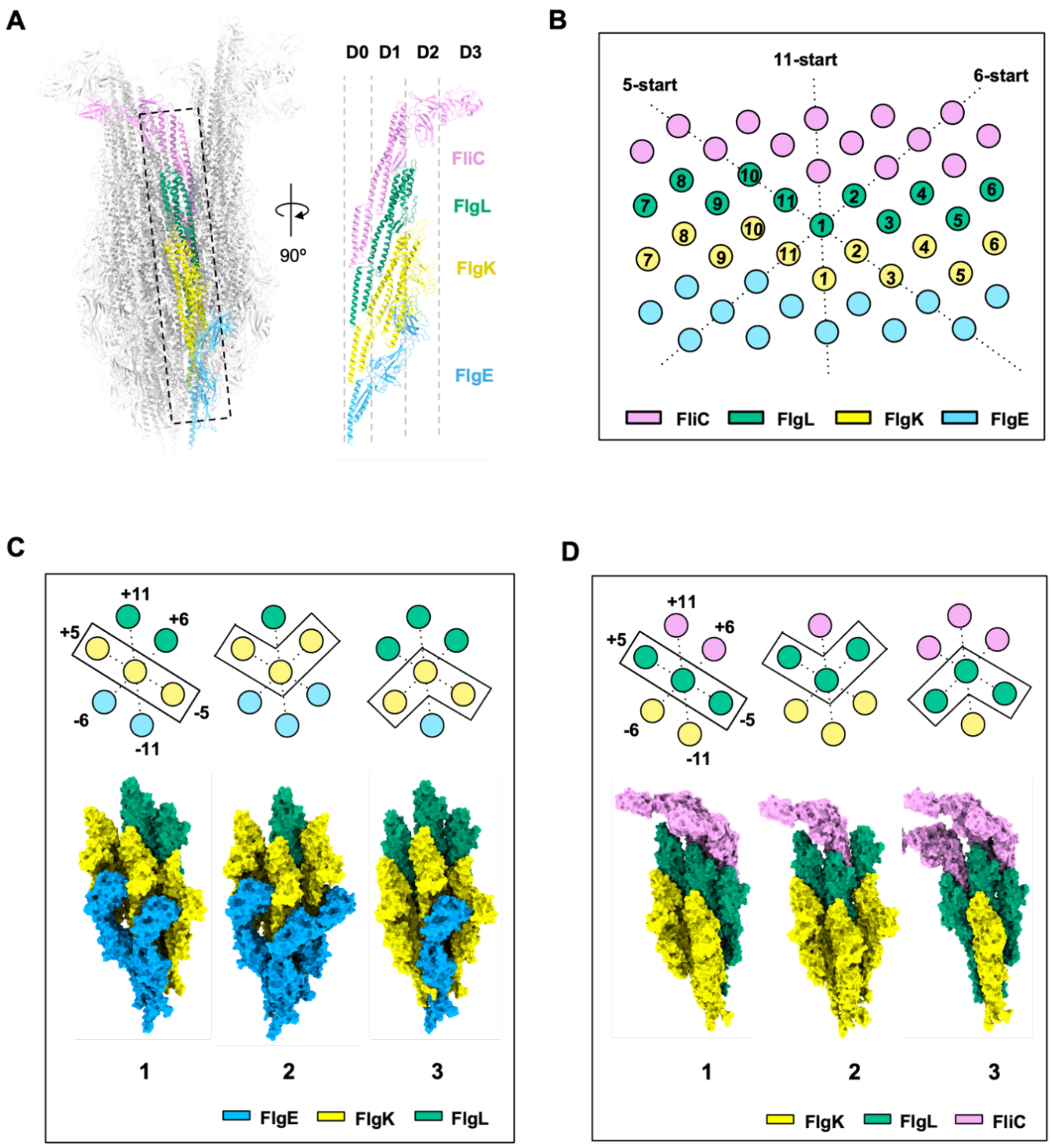


**Figure S5. Details of the interactions of FlgK and FlgL with neighboring subunits, related to Figure 1 and 5.**
(A) Side view and cross section of the protofilament of FlgE-FlgK-FlgL-FliC (PDB: 9GO6).
(B) The arrangement of components in HFJ in the symmetry lattice along 5-start, 6-start, 11-start axes.
(C) FlgK interacts with 6 adjacent proteins in 3 different modes.
(D) FlgL interacts with 6 adjacent proteins in 3 different modes.

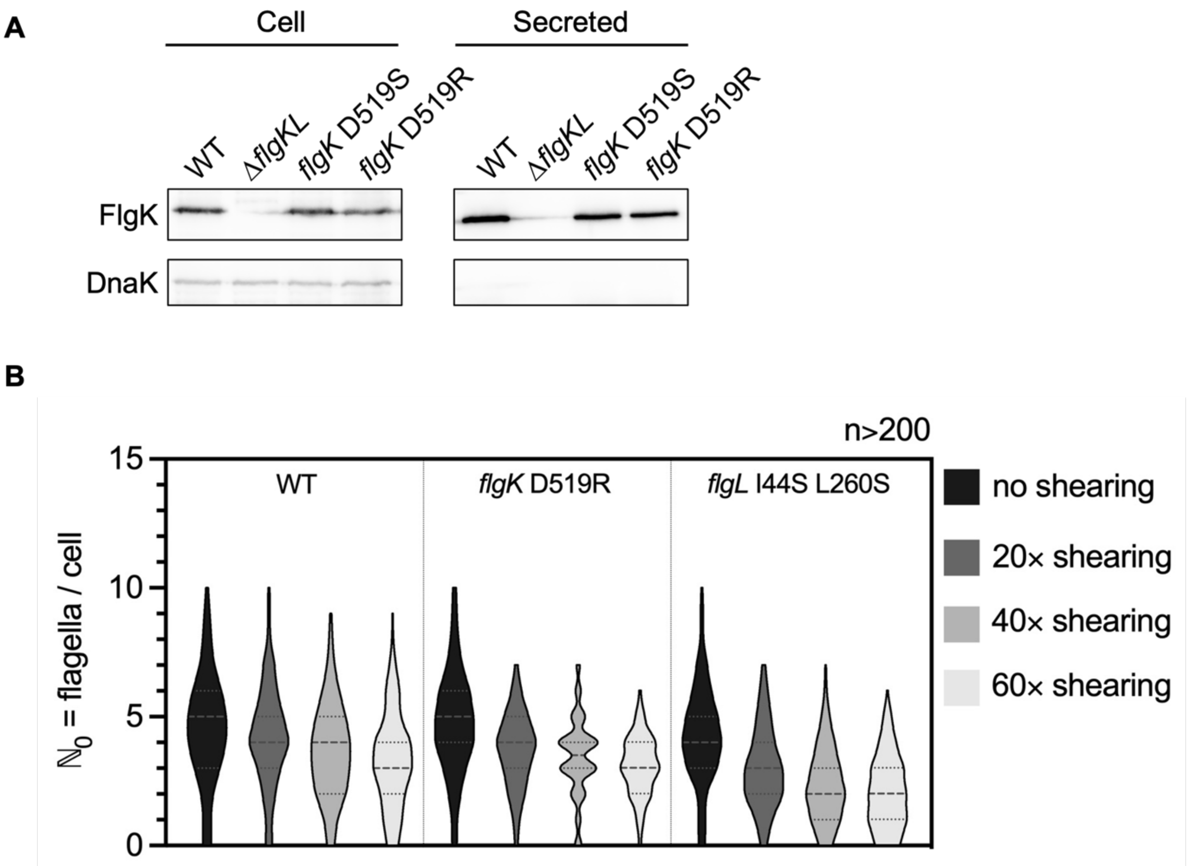


**Figure S6. FlgK mutant variants are secreted in comparable amounts to WT FlgK and flagella of FlgKL interface mutants are more prone to break, related to Figure 5.**
(A) Immunoblotting of the cellular and secreted fractions using the pulsed *flhDC* induction setup of wild-type, a Δ*flgKL* control strain, the FlgK mutants (D519S and D519R) using anti-FlgK antibody. DnaK serves as a loading and lysis control.
(B) Quantification of the number of flagella per cell using the pulsed *flhDC* induction setup in the wild-type and the FlgK D519R and FlgL I44S L260S mutants without shearing and after 20 ×, 40 × or 60 × shearing. Filaments (FliC T237C) were labeled with Dylight555 Maleimide and DNA counterstained with DAPI after pulsed *flhDC* induction. The number of flagella per cell ($\mathbb{N}_0$) was determined for n > 200 individual bacteria per strain and condition for 1 biological replicate. Violin plots represent the distribution of the data including the median (dashed line) and quartiles (dotted lines).

**Supplemental Tables**

**Table S1. Statistics of EM data acquisition, data processing and refinement of atomic models, related to Figures 1 & 6.**

| | FliD cap | Hook-filament junction | Cap-HFJ |
|---|---|---|---|
| **Data collection and processing** | | | |
| Microscope | Thermo Scientific Krios G3i | | Krios |
| Voltage (kV) | 300 | | 300 |
| Detector | K3 | | K2 |
| Magnification | 81,000 | | |
| Pixel size (Å/px) | 1.1 | | 2.2 |
| Dose rate | 16.9 e⁻/pixel/s | | 1.53 e-/Å/frame |
| Total dose (e⁻/Å$^2$) | 43 | | 50 |
| Binning | 2 | | |
| Defocus range (μm) | -0.9 to -2.7 | | -1.5 to -3.0 |
| Total micrographs | 24,729 | | 42,988 |
| Autopicked particles | 527,490 | 5,785,833 | 79,106 |
| Particles after 2D classification | 63,208 | 125,307 | 20,735 |
| Particles in final 3D model | 15,225 | 65,561 | 15,077 |
| Map global resolution (Å) | 3.74 | 2.91 | 6.2 |
| FSC threshold | 0.143 | 0.143 | 0.143 |
| Symmetry imposed | C1 | C1 | C1 |
| EMDB accession | EMD-51486 | EMD-51493 | EMD-51557 |
| **Refinement** | | | |
| Software | PHENIX | PHENIX | PHENIX |
| Model resolution (Å) | 3.73 | 3.17 | 6.5 |
| Map sharpening B factor (Å$^2$) | 23.9 | 18.2 | 18.2 |
| RMSD | | | |
| Bond lengths (Å) | 0.006 | 0.010 | 0.004 |
| Bond angles (°) | 0.668 | 0.890 | 0.833 |
| MolProbity score | 1.84 | 2.56 | 2.47 |
| Clashscore | 12.78 | 17.42 | 26.94 |
| Rotamers outliers (%) | 0.32 | 3.51 | 0.19 |
| Ramachandran plot | | | |
| Favored (%) | 96.53 | 93.99 | 89.90 |
| Allowed (%) | 3.43 | 5.87 | 9.91 |
| Outliers (%) | 0.04 | 0.14 | 0.19 |
| PDB accession | 9GNZ | 9GO6 | 9GSX |

**Table S2. Statistics of the interface area between FlgK or FlgL and their interaction partners, related to Figure 5 and Figure S5.**

| Subject | Mode | Interface area between FlgK/FlgL and adjacent proteins (Å$^2$) | | | | | |
|---|---|---|---|---|---|---|---|
| | | +5 | +11 | +6 | -5 | -11 | -6 |
| FlgK | 1 | 2220.7 (FlgK) | 1688.7 (FlgL) | 456.4 (FlgL) | 2214.9 (FlgK) | 1163.8 (FlgE) | 1045.4 (FlgE) |
| | 2 | 2214.9 (FlgK) | 1568.6 (FlgL) | 992.1 (FlgK) | 1400.5 (FlgE) | 1003.0 (FlgE) | 944.4 (FlgE) |
| | 3 | 1552.9 (FlgL) | 1641.6 (FlgL) | 251.8 (FlgL) | 2188.9 (FlgK) | 1627.6 (FlgE) | 992.1 (FlgK) |
| FlgL | 1 | 1507.2 (FlgL) | 1881.8 (FliC) | 410.6 (FliC) | 1661.9 (FlgL) | 1688.7 (FlgK) | 474.3 (FlgK) |
| | 2 | 1661.9 (FlgL) | 1743.4 (FliC) | 203.6 (FlgL) | 1552.9 (FlgK) | 1568.6 (FlgK) | 456.4 (FlgK) |
| | 3 | 1263.7 (FliC) | 1809.0 (FliC) | 513.5 (FliC) | 1240.4 (FlgL) | 1641.6 (FlgK) | 203.6 (FlgL) |

**Table S3. List of *Salmonella enterica* serovar Typhimurium LT2 strains, related to STAR Methods.**

| Strain | Genotype | Source/Reference |
|---|---|---|
| TH437 | LT2 WT | J. Roth |
| TH9671 | Δ*hin*-5717::FRT (FliC-ON) *fliC*6500(T237C) | Lab collection |
| EM2046 | Δ*hin*-5717::FRT (FliC-ON) *fliC*6500 (T237C) $P_{flhDC}$5451::Tn*10d*Tc[del-25] | Renault et al. [1] |
| EM8327 | Δ*hin*-5717::FRT *flgE*6506 (S171C) $P_{flhDC}$5451::Tn10dTc[del-25] Δ*rflP*::FRT | Lab collection |
| EM8723 | Δ*hin*-5717::FRT Δ*flgKL*5739::FRT *fliC*6500 (T237C) | Lab collection |
| EM8744 | Δ*hin*-5717::FRT *fliC*6500 (T237C) $P_{flhDC}$5451::Tn10dTc[del-25] Δ*flgKL*5739::FRT | Lab collection |
| EM8767 | Δ*fliP*6709 (ΔAA6-241) $P_{flhDC}$5451::Tn10dTc[del-25] Δ*hin*-5717::FRT | Lab collection |
| EM11201 | Δ*hin*-5717::FRT (FliC-ON) *fliC*6500 (T237C) $P_{flhDC}$5451::Tn*10d*Tc[del-25] | This study |
| EM11229 | Δ*hin*-5717::FRT (FliC-ON) *fliC*6500 (T237C) *fliD*23419 (V9S) | This study |
| EM11369 | Δ*hin*-5717::FRT (FliC-ON) *fliC*6500 (T237C) *fliD*23419 (V9S) *fliD*23427(F440S) | This study |
| EM11370 | Δ*hin*-5717::FRT (FliC-ON) *fliC*6500 (T237C) *fliD*23419 (V9S) *fliD*23415 (L443S) | This study |
| EM11371 | Δ*hin*-5717::FRT (FliC-ON) *fliC*6500 (T237C) *fliD*23419 (V9S) *fliD*23416 (M446S) | This study |
| EM11372 | Δ*hin*-5717::FRT (FliC-ON) *fliC*6500 (T237C) *fliD*23419 (V9S) *fliD*23417 (L450S) | This study |
| EM11373 | Δ*hin*-5717::FRT (FliC-ON) *fliC*6500 (T237C) *fliD*23419 (V9S) *fliD*23418 (F461S) | This study |
| EM11374 | Δ*hin*-5717::FRT (FliC-ON) *fliC*6500 (T237C) *fliD*23427(F440S) | This study |
| EM11473 | Δ*hin*-5717::FRT (FliC-ON) *fliC*6500 (T237C) *fliD*23436 (V9R) | This study |
| EM11474 | Δ*hin*-5717::FRT (FliC-ON) *fliC*6500 (T237C) *fliD*23437 (F440R) | This study |

| | | |
|---|---|---|
| EM11475 | Δ*hin*-5717::FRT (FliC-ON) *fliC*6500 (T237C) *fliD*23438 (L443R) | This study |
| EM11476 | Δ*hin*-5717::FRT (FliC-ON) *fliC*6500 (T237C) *fliD*23419 (V9S) *fliD*23437 (F440R) | This study |
| EM11477 | Δ*hin*-5717::FRT (FliC-ON) *fliC*6500 (T237C) *fliD*23419 (V9S) *fliD*23438 (L443R) | This study |
| EM11535 | Δ*hin*-5717::FRT (FliC-ON) *fliC*6500 (T237C) *fliD*23437 (F440R) $P_{flhDC}$5451::Tn*10d*Tc[del-25] | This study |
| EM11727 | Δ*hin*-5717::FRT (FliC-ON) *fliC*6500 (T237C) *fliD*23436 (V9R) $P_{flhDC}$5451::Tn*10d*Tc[del-25] | This study |
| EM11726 | *fliD*23452 (ΔaaS6-F461) $P_{flhDC}$5451::Tn*10d*Tc[del-25] | This study |
| EM15147 | Δ*fliD*23452 (leaves first and last 15 bp) Δ*hin*-5717::FRT | Lab collection |
| EM16009 | P*flhDC*23253::P*proB*-RBS (Δbp -598 to AUG of *flhD*) Δ*hin*-5717::FRT *fliC*5569::*tetRA*(+1 UTR) Δ*sseA*-*ssaU*::FKF (deleted SpiII) | This study |
| EM17200 | Δ*hin*-5717::FRT *fliC*6500 (T237C) *fliD*23823 (L22S) | This study |
| EM17201 | Δ*hin*-5717::FRT *fliC*6500 (T237C) *fliD*23824 (L22R) | This study |
| EM17203 | Δ*hin*-5717::FRT *fliC*6500 (T237C) *fliD*23826 (Y296S) | This study |
| EM17204 | Δ*hin*-5717::FRT *fliC*6500 (T237C) *fliD*23827 (R319S) | This study |
| EM17227 | Δ*hin*-5717::FRT *fliC*6500 (T237C) *flgK*23835 (D519S) | This study |
| EM17234 | Δ*hin*-5717::FRT *fliC*6500 (T237C) *flgK*23837 (Q111S) | This study |
| EM17235 | Δ*hin*-5717::FRT *fliC*6500 (T237C) *flgK*23838 (Q118S) | This study |
| EM17237 | Δ*hin*-5717::FRT *fliC*6500 (T237C) *flgL*23840 (I44S) | This study |
| EM17241 | Δ*hin*-5717::FRT *fliC*6500 (T237C) *flgL*23844 (L260S) | This study |
| EM17307 | Δ*hin*-5717::FRT *fliC*6500 (T237C) *fliD*23853 (Y296S, R319S) | This study |
| EM17316 | Δ*hin*-5717::FRT *fliC*6500 (T237C) *flgK*23854 (Q111S, Q118S) | This study |
| EM17330 | Δ*hin*-5717::FRT *fliC*6500 (T237C) *flgK*23835 (D519S) P*flhDC*5451::Tn10dTc[del-25] | This study |
| EM17353 | Δ*hin*-5717::FRT *fliC*6500 (T237C) *fliD*23858 (Y296R) | This study |

| | | |
|---|---|---|
| EM17355 | Δ*hin*-5717::FRT *fliC*6500 (T237C) *flgK*23860 (D519R) | This study |
| EM17356 | Δ*hin*-5717::FRT *fliC*6500 (T237C) *flgK*23861 (Q111S, D519S) | This study |
| EM17357 | Δ*hin*-5717::FRT *fliC*6500 (T237C) *flgK*23862 (Q118S, D519S) | This study |
| EM17358 | Δ*hin*-5717::FRT *fliC*6500 (T237C) *flgL*23863 (I44R) | This study |
| EM17360 | Δ*hin*-5717::FRT *fliC*6500 (T237C) *flgL*23865 (L260R) | This study |
| EM17361 | Δ*hin*-5717::FRT *fliC*6500 (T237C) *flgL*23866 (I44S, L260S) | This study |
| EM17362 | Δ*hin*-5717::FRT *fliC*6500 (T237C) *flgK*23835 (D519S) *flgL*23840 (I44S) | This study |
| EM17390 | Δ*hin*-5717::FRT *fliC*6500 (T237C) *flgK*23860 (D519R) P*flhDC*5451::Tn10dTc[del-25] | This study |
| EM17399 | Δ*hin*-5717::FRT *fliC*6500 (T237C) *flgL*23866 (I44S, L260S) P*flhDC*5451::Tn10dTc[del-25] | This study |

**Table S4. List of oligonucleotides used in this study, related to STAR Methods.**

| Sequence | Source | Identifier |
|---|---|---|
| CAAAAAGGAAGAAGGCATGGCTTCAATTTCAT CATTAGGTAGGGTTTTCCCAGTCACGAC | This study | fliD_V9::Kan_fw |
| TCAGGTCTGTCAACAACTGGTCTAACGGTAAG TTTGATCCTGCTTCCGGCTCGTATGTTG | This study | fliD_V9::Kan_rv |
| CAAGGCCCAGTTTACCCAACTGGATACCATGA TGAGTAAGAGGGTTTTCCCAGTCACGAC | This study | fliD_L450_F641::Kan_fw |
| TACATGGTGACCTCTGTTATCAGGACTTGTTC ATAGCTGTTGCTTCCGGCTCGTATGTTG | This study | fliD_L450_F641::Kan_rv |
| TGCAATCAAAAAGGAAGAAGGCATGGCTTCAA TTTCATCAAGGGTTTTCCCAGTCACGAC | This study | fliD_L7-L22::Kan/SceI_fw |
| GTTTGGTAATTGGCGTTAAGCGTCCTTTTTCG TTCTTTGTTGCTTCCGGCTCGTATGTTG | This study | fliD_L7-L22::Kan/SceI_rv |
| CTATAACTCGCTGGTGGATACCTTTAGCTCGT TAACCAAAAGGGTTTTCCCAGTCACGAC | This study | fliD-KanSceI Daa296-319 fw |
| CGCTATTGGCAAATTGTGCCCGAATCCCGGTC TGGATAGTTGCTTCCGGCTCGTATGTTG | This study | fliD-KanSceI Daa296-319 rev |
| CCTGCTGGCCGATAAATCCAGCTCACTGTCTG GTTCGTTGAGGGTTTTCCCAGTCACGAC | This study | flgK-KanSce Daa111-118 fw |
| CCTGACGCGCCGCAGGATCTTCCGCGTTACT GACTAACGTTGCTTCCGGCTCGTATGTTG | This study | flgK-KanSce Daa111-118 rev |
| GATTGCGATGGCGTCTGAGTCAAAACTCGATC CTGACGTGAGGGTTTTCCCAGTCACGAC | This study | flgK-KanSceI Daa447-518 fw |
| AATACTGCTGATAACGCTGCAAATTGCCGTAC TCTTCGTCTGCTTCCGGCTCGTATGTTG | This study | flgK-KanSceI Daa447-518 rev |
| CCAACCCATCTGACGATCCGATCGCCGCGTC GCAGGCGAGGGTTTTCCCAGTCACGAC | This study | flgL-KanSceI Daa50-139 fw |
| CCACCTGTCGCCTGGTCGAATGGCGCCGCTT CCGTTTTTGCTTCCGGCTCGTATGTTG | This study | flgL-KanSceI Daa50-139 rev |
| CGTGCGGAACTGGGAACGCAACTGAGCGAAC TCAGTACGAGGGTTTTCCCAGTCACGAC | This study | flgL-KanSceI Daa260-304 fw |
| GTTACCGGTTCAACTGGAAAAGCGACATTCCC TGCATGTCTGCTTCCGGCTCGTATGTTG | This study | flgL-KanSceI Daa260-304 rev |
| AAAGGAAGAAGGCATGGCTTCAATTTCATCAT TAGGTAGCGGATCAAACTTACCGTTAGA | This study | fliD_V9S_fw |
| AAAGGAAGAAGGCATGGCTTCAATTTCATCAT TAGGTCGTGGATCAAACTTACCGTTAGA | This study | fliD_V9R_fw |
| TTTGGTAATTGGCGTTAAGCGTCCTTTTTCGTT CTTTGTGCTGTCTGTCAACAACTGGTC | This study | fliD L22S fw NEW |
| TTTGGTAATTGGCGTTAAGCGTCCTTTTTCGTT CTTTGTACGGTCTGTCAACAACTGGTC | This study | fliD L22S rev NEW |
| TATAACTCGCTGGTGGATACCTTTAGCTCGTT AACCAAAAGCACCGCCGTTGAGCCGGGC | This study | fliD Y296S fw |
| TATAACTCGCTGGTGGATACCTTTAGCTCGTT AACCAAACGCACCGCCGTTGAGCCGGGC | This study | fliD Y296R |
| CTATTGGCAAATTGTGCCCGAATCCCGGTCTG GATAGTGCTAACCACACTATCGCCTAAC | This study | fliD R319S rev |

| Sequence | Source | Identifier |
|---|---|---|
| CAGCATCGATGAAACCGTTGCCCGTTACAAG GCCCAGAGCACCCAACTGGATACCATGAT | This study | fliD_F440S_fw |
| CAGCATCGATGAAACCGTTGCCCGTTACAAG GCCCAGCGTACCCAACTGGATACCATGAT | This study | fliD_F440R_fw |
| TGAAACCGTTGCCCGTTACAAGGCCCAGTTTA CCCAAAGCGATACCATGATGAGTAAGCT | This study | fliD_L443S_fw |
| TGAAACCGTTGCCCGTTACAAGGCCCAGTTTA CCCAACGTGATACCATGATGAGTAAGCT | This study | fliD_L443R_fw |
| GGCCCAGTTTACCCAACTGGATACCATGATGA GTAAGAGCAATAACACCAGTAGTTATTT | This study | fliD_L450S_fw |
| TGCCCGTTACAAGGCCCAGTTTACCCAACTGG ATACCAGCATGAGTAAGCTGAATAACAC | This study | fliD_M446S_fw |
| ATGGTGACCTCTGTTATCAGGACTTGTTCATA GCTGTGCTTTGCTGGGTCAAATAAC | This study | fliD_F461S_rv |
| CTGCTGGTAGGCGATGGTAA | This study | fliD_fr1 |
| GCCAGCGGTAGTACTGACTT | This study | fliD_rv1 |
| GCTTTCAGCCCGTTATCGAT | This study | fliD_rv2 |
| TCAATCAACTGATGCGGGCT | This study | fliD_3UTR_rv |
| TCTTAGGCGGCGAAATAGCC | This study | fliD_5UTR_fw |
| GATCACGGTGGAAGGCGATA | This study | fliD_internal_fw |
| CGCGTTTTCTGCTTTCACCA | This study | fliD_internal_rv |
| CTGCTGGCCGATAAATCCAGCTCACTGTCTGG TTCGTTGAGCAGTTTTTTTACCAGCCTG | This study | flgK Q111S fw |
| CTGACGCGCCGCAGGATCTTCCGCGTTACTG ACTAACGTGCTCAGGCTGGTAAAAAAACT | This study | flgK Q118S rev |
| ATAATACTGCTGATAACGCTGCAAATTGCCGT ACTCTTCGCTGAGGTTAACGCCGGAAAC | This study | flgK D519S rev |
| ATAATACTGCTGATAACGCTGCAAATTGCCGT ACTCTTCGCGGAGGTTAACGCCGGAAAC | This study | flgK D519R rev |
| CTACCGGTAAGCGCGTTACCAACCCATCTGAC GATCCGAGCGCCGCGTCGCAGGCGGTAG | This study | flgL I44S fw |
| CTACCGGTAAGCGCGTTACCAACCCATCTGAC GATCCGCGCGCCGCGTCGCAGGCGGTAG | This study | flgL I44R fw |
| GTGCGGAACTGGGAACGCAACTGAGCGAACT CAGTACGAGCGATTCACTGGGAAGCGAC | This study | flgL L260S fw |
| GTGCGGAACTGGGAACGCAACTGAGCGAACT CAGTACGCGCGATTCACTGGGAAGCGAC | This study | flgL L260R fw |
| ACTGACCCGAATTACGCGC | Lab Collection | flgK_sequ_fw_53C |
| GCTTACCGGTAGACATCTGC | Lab Collection | flgK_sequ_rv_54C |

| Sequence | Source | Identifier |
|---|---|---|
| CGGGAGATTTGATCGTTCAGG | Lab Collection | FlgK_int_seq_rv |
| TCGACAGCACGAAGGTTCAG | Lab Collection | 5'_flgK_int1077_seq_fw |
| AGAACGGAACGTCAGTAGCC | Lab Collection | 3'_flgK_int844_seq_rv |
| CAGTCATAGCCGAATAGCCT | Lab Collection | K1-test |
| GGCGTTAACCTCGACGAAGA | Lab Collection | 5'_flgK_seq_fw |
| CTACGGGCGCGACAATATGA | Lab Collection | 3'_Int_flgL_rne_seq_rv |

**Supplemental Videos**

**Movie S1. Cryo-ET series of the native flagellar filament cap complex of intact *S. enterica* cells, related to Figure S1.**

**Movie S2. The overview of the extracellular flagella of *S. enterica*, related to Figure 1.**
Density of hook, junction, filament and cap are segmented and colored as in Figure 1D. The movie shows a 360° view of the overall structure, focus views of each section and the atomic model that is built from the density map.

**Movie S3. Visualizing FliD movement upon FliC incorporation by 3D variability analysis at the front and the top view, related to Figure 3.**

**Movie S4. The rise and rotation of FliD, related to Figure 3.**
The central FliD in the motions is colored in red while the FliD before and after along the CCW direction are colored in salmon and orange. FliDs that are unmoved during the rotation are colored in white.

**Movie S5. Focused view on D0-D1 domains during the FliD movement in Movie S4, related to Figure 3.**

**Movie S6. The morphing from *C. jejuni* FliD pentamer structure to *S. enterica* FliD pentamer structure, related to Figure 6.**
The FliD with significant conformational changes is colored in red.
